# An Evolutionarily Conserved Ypk1/SGK1 Kinase Pathway Regulates the Unfolded Protein Response by Modulating the Ire1 Protein Abundance

**DOI:** 10.64898/2026.09.11.751087

**Authors:** Saswata Chakrabarty, Anish Chakraborty, Jagadeesh Kumar Uppala, Noelle Marie Bryan, Nadège Gouignard, Nicholas J Reiter, An Phu Tran Nguyen, Madhusudan Dey

## Abstract

The unfolded protein response (UPR) is a cellular mechanism that maintains protein homeostasis (proteostasis) under conditions of endoplasmic reticulum (ER) stress. The dual kinase/RNase Ire1 is a conserved regulator of the UPR, mediating the unconventional cytosolic splicing of *HAC1* mRNA in yeast and *XBP1* mRNA in human cells. The resulting spliced *HAC1/XBP1* transcript encodes a transcription factor that induces the expression of protein-folding chaperones and stress-responsive genes, thereby restoring proteostasis. In our previous work, we showed that the MAP kinase Slt2 (homolog of human ERKs) contributes to UPR signaling by promoting *IRE1* expression through the transcription factor Rlm1 (homolog of human MEF2C). Here, we demonstrate that Hac1 expression is reduced in yeast strains deficient in essential protein kinase Cdc28, Pkc1, Rio2, Tor2, Pkh1, or Ypk1, suggesting that these kinases also serve as UPR regulators. We focused on the kinase Ypk1, the yeast ortholog of human SGK1 (serum/glucocorticoid-regulated kinase 1). We provide genetic and biochemical evidence that Ypk1/SGK1 acts upstream of the Pkc1/PKCδ signaling pathway and is required for maintaining IRE1 protein abundance in both yeast and human cells. Collectively, our results identify an evolutionarily conserved Ypk1/SGK1 signaling pathway that regulates the *HAC1* and XBP1 mRNA splicing by modulating the Ire1 protein abundance.

**SIGNIFICANCE STATEMENT:** Ypk1, the yeast ortholog of human SGK1, and Ire1, the yeast ortholog of human IRE1, are key regulators of lipid and protein homeostasis, respectively. Here, we provide genetic and biochemical evidence that the Ypk1/SGK1 signaling pathway promotes protein homeostasis by controlling Ire1/IRE1 protein abundance.

This fundamental study has broader implications for the field of protein kinases and drug discovery, as both IRE1 and SGK1 are attractive therapeutic targets for a range of diseases, including blood diseases (PMC5338400), aging (PMC9670206) and certain cancers (PMC7851074). Many compounds are developed to modulate the IRE1 or SGK1 activity, but their clinical use remains limited, due in part to their off-target effects and incomplete understanding of their regulation and biology. Thus, identifying novel crosstalk and co-regulation between IRE1 and SGK1 signaling pathways will provide new opportunities to selectively modulate IRE1 and/or SGK1 activity and facilitate the development of more effective therapeutics.

## INTRODUCTION

The endoplasmic reticulum is a cellular compartment where most secretory proteins fold and mature to become biologically active forms(1). Additionally, ER plays a significant role in Ca^2+^ homeostasis and lipid biosynthesis(2,3). Hence, any perturbation in ER functions leads to the accumulation of unfolded or mis-folded proteins inside the ER, a condition known as the “ER stress”. ER stress evokes adaptive or apoptotic ER stress response (ESR) depending on the type, severity, and duration of stress. Two major ESRs are the unfolded protein response (UPR) (4–7) and the heat shock response (HSR) (8). The UPR initiates a network of signaling pathways, which dynamically reprogram the cellular physiology to mitigate the ER stress. These include (1) attenuation of translation of certain mRNAs, (2) enhanced expression of protein folding enzymes and chaperones, (3) activation of the ER-associated degradation (ERAD) of unfolded proteins, and (4) initiation of the apoptotic program when ER stress is irremediable.

In metazoan cells, the UPR is initiated by three major sensors: IRE1 (inositol-requiring enzyme 1)(9–11), PERK (protein kinase RNA-like ER kinase) (12), and ATF6 (activating transcription factor-6) (13). IRE1 is the conserved sensor and is the sole ER stress sensor in the budding yeast *Saccharomyces cerevisiae*(14–16), which contains an N-terminal lumenal domain (LD) and C-terminal cytosolic domain (Ire1^cyto^). Ire1^cyto^ has a kinase domain (KD) and an RNase domain known as KEN (kinase extension nuclease) domain. The KEN domain initiates the UPR signal by cleaving an inhibitory intron from the *HAC1* mRNA in yeast cells (17–20) or *XBP1* mRNA in metazoan cells (21,22). The cleaved mRNAs are then ligated by tRNA ligase in yeast cells (23) and RTCB in mammalian cells (24). The spliced *HAC1*/*XBP1* mRNA translates a transcription factor that activates the expression of protein-folding enzymes and chaperones that, in turn, enhance the ER proteostasis.

RNA microarray analysis in yeast cells shows that Hac1 protein pleiotropically activates approximately 430 mRNAs in response to ER stress (25). Like Hac1 in yeast cells, XBP1 and ATF4 in human cells are reported to activate numerous mRNAs in response to ER stress (26,27). Encoded proteins from these mRNAs include molecular chaperones, protein disulfide isomerase, and thiol oxidase (25,28). Dalfsen et al. (2018) showed that a small subset of Hac1-driven genes, specifically genes encoding proteins for aerobic respiration, are translationally down-regulated by their long un-decoded transcript isoforms (LUTIs) (29). Of note, only a subset of those mRNAs is shown to be involved in enhancing the protein-folding capacity of ER, and the roles of many transcriptionally activated genes are still unknown.

Multiple protein kinases play regulatory roles in yeast UPR by as-yet unknow mechanisms. The mitogen-activated protein (MAP) kinase Slt2, ortholog human ERKs, plays a regulatory role in the ER stress response (30,31). We also show that the splicing of *HAC1* pre-mRNA is partially regulated by protein kinases Kin1 and Kin2 (27), whereas the translation of matured *HAC1* mRNA is regulated by Vps34 and Tor kinases (32). Protein kinases Pkh1/2, orthologues of human PDK1 (3-phosphoinositide-dependent kinase 1), activate the ER-stress-induced transcriptional response (33). Pkh1/2 is known to phosphorylate the protein kinase Ypk1/2 [orthologs of human serum/glucocorticoid-regulated kinase 1(SGK1)] that controls cellular endocytosis via sphingolipid-mediated signaling pathway (34,35). Together, these studies indicate the complexity of the ER stress response, which is facilitated by the concerted activity of multiple kinases. In this study, we show that protein kinases Cdc28, Pkc1, Rio2, Tor2, Pkh1/2, and Ypk1/2 are UPR regulators in yeast cells. We focused on the kinase Ypk1, the yeast ortholog of human SGK1, and provide genetic and biochemical evidence for a novel, evolutionarily conserved mechanism for UPR in which Ypk1/SGK1 regulates the *HAC1/XBP1* mRNA splicing by regulating the IRE1 protein abundance.

## RESULTS

### 1. Essential protein kinases Cdc28, Pkc1, Rio2, Tor2, Pkh1/2 and Ypk1/2 contribute to ER stress response

To investigate the contribution of essential kinases to the ER stress response, we examined the Hac1 protein levels in 17 yeast *S. cerevisiae* strains carrying temperature-sensitive (ts)-mutations in essential kinases after a short incubation at non-permissible temperature. First, we analyzed growth of ts-strains at permissive temperature (25°C) and non-permissive temperatures (37°C or 38°C) on a solid YEPD medium (**Fig 1A**). Thirteen strains harbored a ts-mutation of the corresponding kinase gene (36). The remaining 4 strains carried a ts-mutation of the targeted kinase and a deletion of its paralog such as *tor1Δtor2^ts^*(37), *pkh1^ts^pkh2Δ*(38), *ypk1^ts^ypk2Δ*(39) and *yck1^ts^yck2Δ* (40). At 25°C, all strains grew on YEPD medium, although some exhibited slower growth (**Fig 1A**, left panel). In contrast, growth was retarded or completely abolished at 37°C (**Fig 1A**, right panel and **Supplemental Fig S1**). Notably, when culture was incubated at 37°C for 3 hours and subsequently returned to 25°C, growth was restored, indicating that ts-mutations are not lethal, rather the activity of ts-allele is reversibly impaired at non-permissive temperature and recovered at permissive temperature.

**Fig 1:**
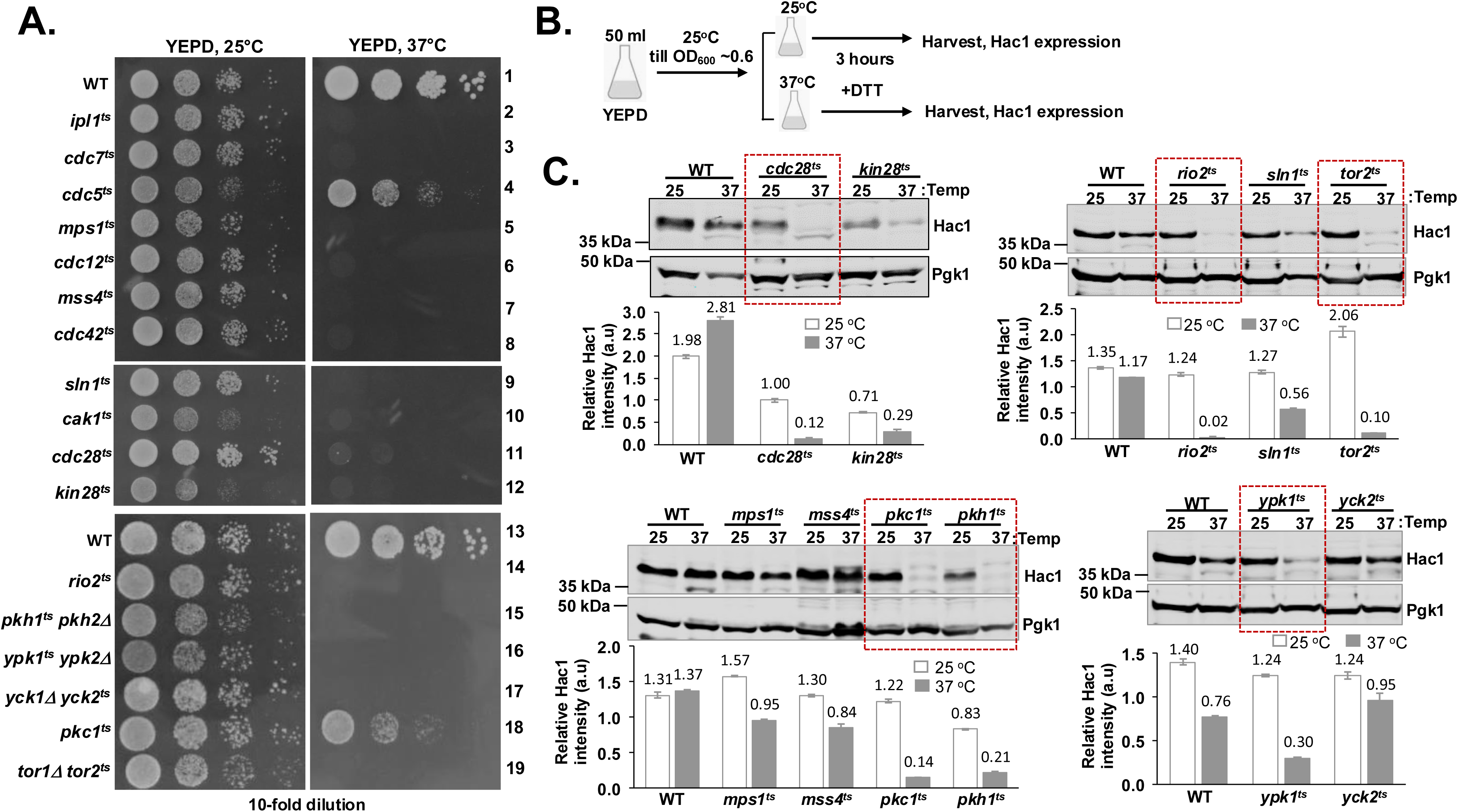
Analysis of Hac1 protein expression in temperature-sensitive yeast strains. **(A)** Indicated yeast strains were serially diluted, spotted on the YEPD medium and grown at 25°C and 37°C for 48 hours. **(B)** The schema provides a detailed representation of the sequential steps involved in growing cells, stress induction by DTT for three hours, and harvesting the cultures. **(C)** Whole cell extracts (WCEs) were prepared from the indicated strains at 25°C and 37°C and subjected to Western blot analysis by Hac1 and Pgk1 antibodies. The intensities of Hac1 and Pgk1 protein bands were measured, and the relative intensities as arbitrary units (a. u.) are shown as a bar diagram at the bottom.

The ts-strains were then cultured in a liquid YEPD medium at both 25°C and 37°C or 38°C (only for the *pkc1^ts^* strain) for 3 hours in the presence of an ER stressor DTT (**Fig 1B)** and monitored for their ability to express Hac1 protein. At 25°C, Hac1 protein expressions in all ts-strains were almost comparable with WT (**Fig 1C** and **Supplemental Fig S1)**. In contrast, the Hac1 expressions were markedly reduced in the *cdc28^ts^, pkc1^ts^, pkh1^ts^pkh2Δ, rio2^ts^, tor1Δtor2^ts^, and ypk1^ts^ypk2Δ* strains at 37°C (**Fig 1C**). These results suggest that protein kinases Cdc28, Pkc1, Pkh1, Rio2, Tor2 and Ypk1 independently or coordinately contribute to the Hac1-mediated ER stress response in addition to their essential functions, such as the cell cycle regulation by Cdc28(41) and Tor2 (42), the cell wall remodeling by Pkc1(43), the ribosome biogenesis by Rio2(44), and the lipid metabolism by Pkh1 and Ypk1(45,46).

We primarily focused here on the kinase Ypk1/2 and it’s human ortholog SGK1 with some additional work on yeast kinases Pkh1 and Pkc1. Ypk1/2 is known to be phosphorylated and activated by two kinases: Tor2 complex (47,48) and Pkh1/2 (39). Once activated, Ypk1/2 phosphorylates and activates several downstream targets, including the protein kinase Fpk1/2(49) and dehydrogenase Gpd1 (glycerol-3-phosphate dehydrogenase)(50) that regulate lipid metabolism, as well as the ER trans-membrane protein Orm1/2 that is reported to maintain the protein quality control (51). Thus, involvement of Ypk1 signaling in Hac1-mediated ER stress response identifies a novel and uncharacterized role for this pathway.

Protein kinases Ypk1/2, Pkh1/2, and Pkc1 are orthologs of human kinases SGK1, PDK1, and novel PKC (nPKC) isoforms nPKCδ and nPKCη(52). Sequence analysis showed that, in addition to their conserved kinase domains, they contain several distinct protein domains. Ypk1/2 possesses a unique N-terminal C2 domain that is absent in human SGK1**(Supplemental Fig S2**). Both Pkh1/2 and its human PDK1 contain a C-terminal pleckstrin homology (PH) domain **(Supplemental Fig S3**). Pkc1 harbors unique HR1 and C2 domains that are absent in human PKCδ/PKCη **(Supplemental Fig S4**). Consistent with previous reports (34,52), expression of these human kinases SGK1, PDK1 and the PKCδ/PKCη complemented the ts-phenotype of the respective yeast kinases (**Supplemental Figs. S2, S3** and **S4**), confirming their functional conservation of these signaling proteins between yeast and human cells **(Supplemental Fig S5**).

### 2. Reduced Hac1 expression in the *pkh1^ts^*, *ypk1^ts^* or *pkc1^ts^* strain is caused by diminished *HAC1* mRNA splicing

To confirm that Ypk1/2, Pkh1/2 and Pkc1 contribute to Hac1-mediated UPR, we also compared Hac1 protein expression in WT, *pkh1^ts^pkh2Δ, ypk1^ts^ypk2Δ* and *pkc1^ts^* strains at permissive (25°C), semi-permissive (30°C) and non-permissive (37°C or 38°C for *pkc1^ts^*) temperatures. In WT cells, Hac1 protein was expressed when cells were grown in the presence of DTT at 25°C, 30°C, and 37°C (**Fig 2A**). Compared to cells grown at 25°C, Hac1 protein expression was reduced in *pkh1^ts^pkh2Δ* (∼1.5-fold at 30°C and ∼8-fold at 37°C), *ypk1^ts^ypk2Δ* (∼1.5-fold at 30°C and ∼10-fold at 37°C), *pkc1^ts^* (∼1.6-fold at 30°C and ∼4.6-fold at 38°C) strains (**Fig 2A**). These reductions in Hac1 protein expression in the ts-strains further confirm that kinases Ypk1/2, Pkh1/2 and Pkc1 contribute to Hac1-mediated ER stress response.

**Fig 2:**
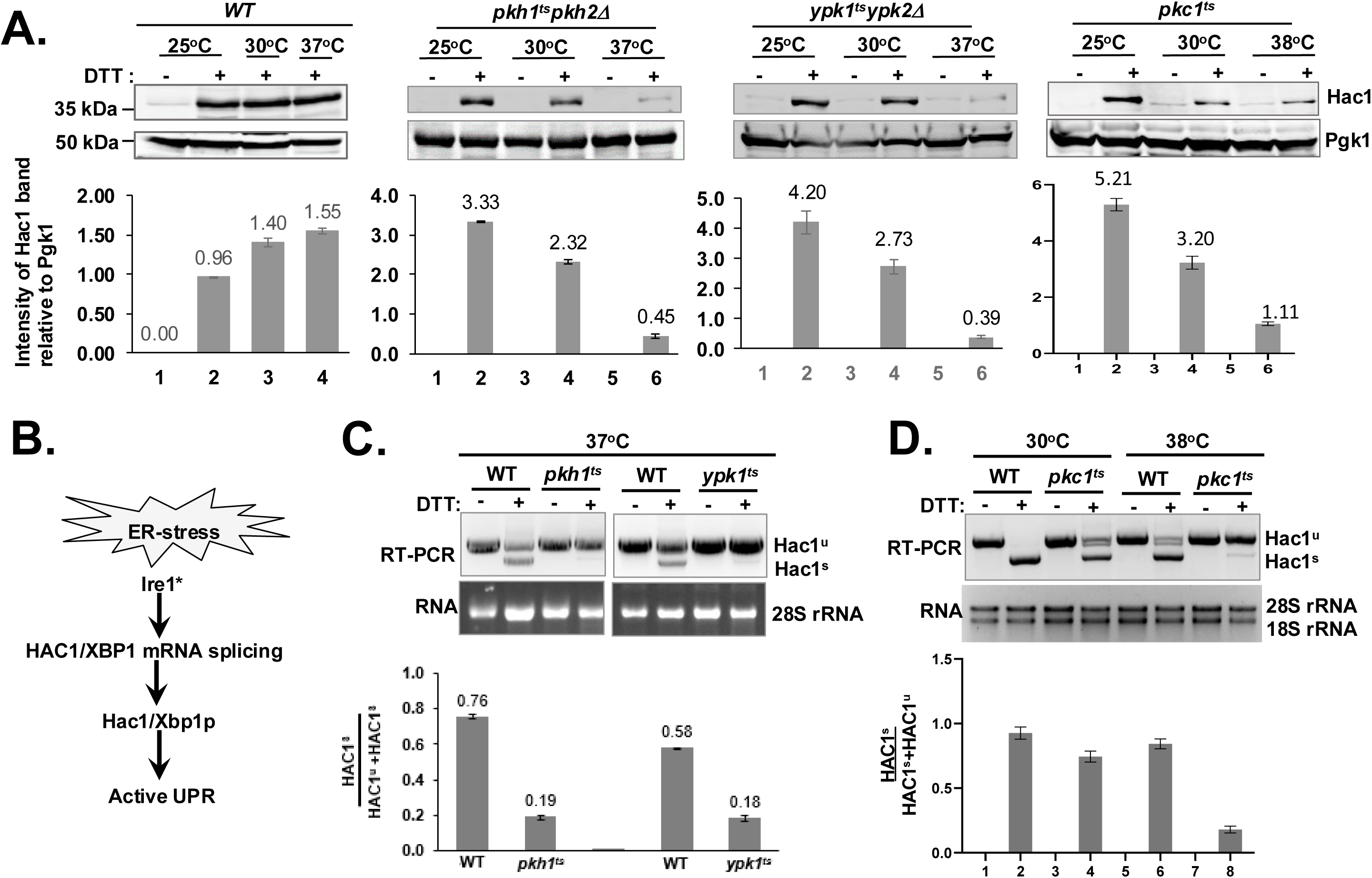
Reduced *HAC1* mRNA splicing in Pkh1, Ypk1 and Pkc1 deficient cells. **(A)** The indicated yeast strains were grown at 25°C and 37°C (38°C for the *pkc1^ts^* strain) in the presence and absence of DTT. WCEs were prepared and subjected to Western blot analysis using Hac1 and Pgk1 antibodies. The intensities of Hac1 and Pgk1 protein bands were measured were measured, and the relative intensities of Hac1 protein bands were shown as a bar diagram at the bottom. **(B)** The schematic representation of Ire1-Hac1/XBP1 pathway. (**C & D**) The indicated WT and its isogenic ts-strains were grown at 25°C and 37°C (38°C for the *pkc1^ts^* strain) in the presence and absence of DTT. Total RNA was isolated and subjected to RT-PCR analysis to monitor un-spliced (HAC1^u^) and spliced (HAC1^s^) forms of *HAC1* mRNA.

Loss of both Pkh1 and Pkh2 or Ypk1 and Ypk2 isoforms reduced the Hac1 expression (**Fig 2A**); however, yeast cells lacking only an individual kinase, whether it be Pkh1, Pkh2, Ypk1, or Ypk2, grew on the medium contain an ER stressor tunicamycin comparable to that of the WT strain and produced almost similar levels of Hac1 protein during an ER stress (**Supplemental Fig S6**), supporting the notion that isoforms perform similar function. This redundancy is likely a robustness of mechanisms, allowing cells to have higher adaptive capacity to adverse and diverse environmental conditions.

Next, we investigated if the reduced level of Hac1 protein in the *pkh1^ts^pkh2Δ*, *ypk1^ts^ypk2Δ* or *pkc1^ts^* strain was due to inefficient cytosolic splicing of *HAC1* pre-mRNA (**Fig 2B**). WT, *pkh1^ts^pkh2Δ* and *ypk1^ts^ypk2Δ* strains were grown at 37°C in the presence and absence of DTT for three hours, whereas the *pkc1^ts^* strain was grown at both 30°C and 38°C under the same conditions. Total RNAs were isolated and the levels of spliced (HAC1^s^) and un-spliced (HAC1^u^) forms of *HAC1* mRNA were compared by RT-PCR. *HAC1^u^* mRNA was detected in WT, *pkh1^ts^pkh2Δ*, *ypk1^ts^ypk2Δ* and *pkc1^ts^* cells when grown with or without DTT (**Figs 2C and 2D**). Both *HAC1^s^* and *HAC1^u^* mRNA species were observed in only WT cells when grown in the presence of DTT (**Fig 2C**). In contrast, *HAC1^s^* mRNA species were significantly reduced in *pkh1^ts^pkh2Δ* (**Fig 2C)**, *ypk1^ts^ypk2Δ* (**Fig 2C)** and *pkc1^ts^* strains (**Fig 2D**). The reduced cytosolic splicing of *HAC1* mRNA suggests that protein kinases Pkh1, Ypk1 and Pkc1 are functionally linked to Ire1-mediated ER stress response.

### 3. Reduced *HAC1* mRNA splicing in *pkh1^ts^*, *ypk1^ts^* or *pkc1^ts^* strain is caused by reduced Ire1 protein abundance

To determine whether the reduced level of *HAC1* mRNA splicing observed in the *pkh1^ts^pkh2Δ*, *ypk1^ts^ypk2Δ* and *pkc1^ts^* strain was caused by decreased Ire1 expression or impaired Ire1 activation **(Fig 3A)**, we examined the Ire1 protein levels in these strains at both 25°C and 37°C or 38°C (*pkc1^ts^* strain). The endogenous expression of Ire1 protein is low and nearly undetectable by Western blot analysis **(Fig 3B)**. Therefore, to assess Ire1 protein levels, we introduced a plasmid (pIRE1) expressing Ire1 under its native promoter (pIRE1) in these strains. WT cells containing the plasmid pIRE1, the Ire1 protein level was increased after 2 and 4 hours of DTT treatment (**Fig 3B**), suggesting that Ire1 expression was induced under an ER stress.

**Fig 3:**
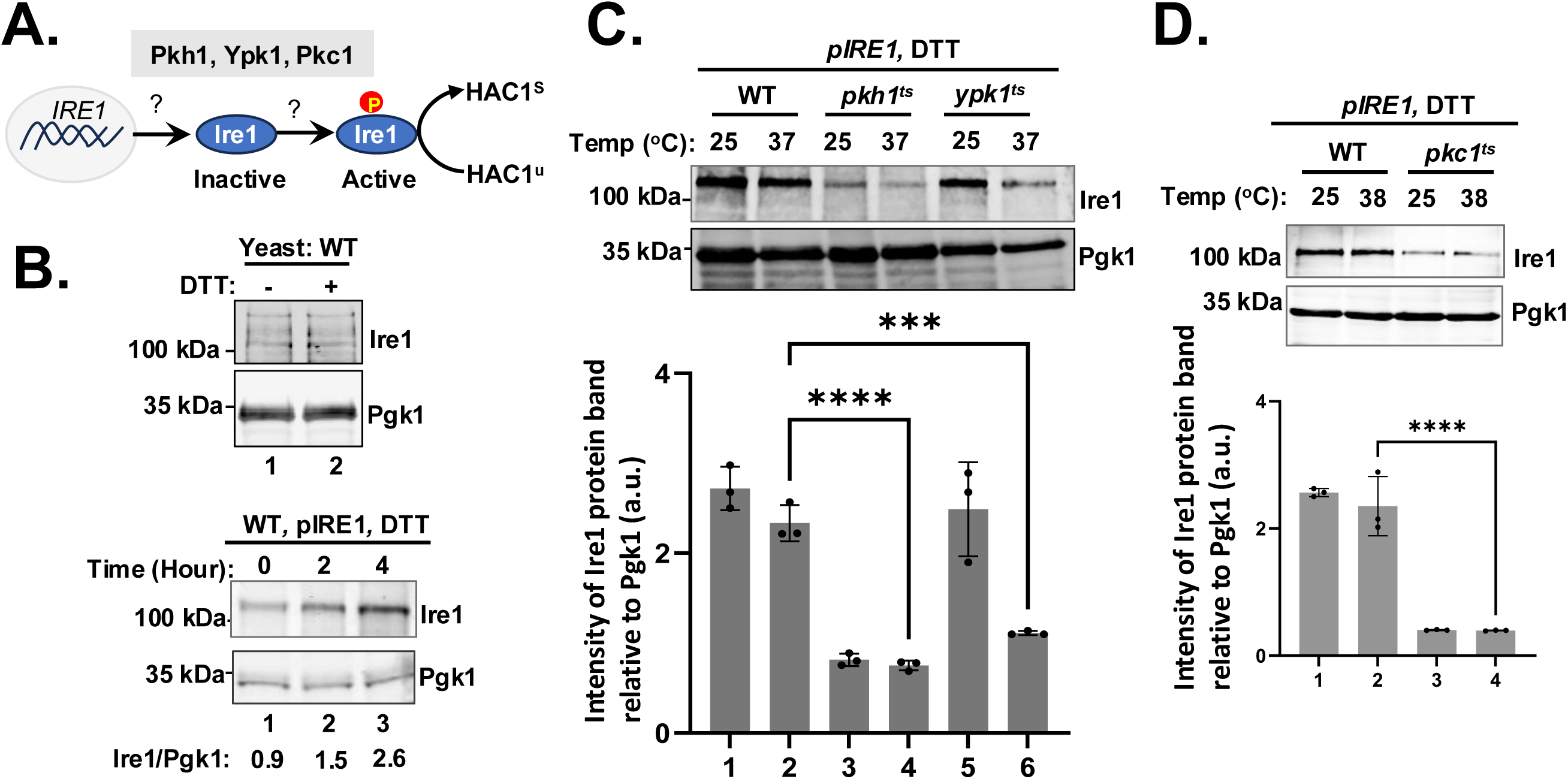
Reduced Ire1 protein abundance in Pkh1, Ypk1 and Pkc1 deficient cells. **(A)** The schematic representation of *IRE1* gene expression, Ire1 protein activation and *HAC1* mRNA splicing. **(B)** (Upper panels) Wild type yeast strain was grown in YEPD medium in the presence and absence of DTT. WCEs were prepared and subjected to Western blot analysis using Ire1 and Pgk1 antibodies. (Lower panels) Wild type yeast strains expressing Ire1 from a 2µ plasmid from its native promoter (pIRE1) was grown in SC-uracil medium in the presence of DTT. WCEs were prepared and subjected to Western blot analysis using Ire1 and Pgk1 antibodies. **(C)** & (**D**) Wild type and the indicated ts-strains were transformed with the plasmid pIRE1. Transformants were grown in the presence of DTT. WCEs were prepared and subjected to Western blot analysis using Ire1 and Pgk1 antibodies. Experiments were repeated at least three times. The intensities of protein bands were measured and shown in a bar diagram (****p-value<0.0001, paired t-test).

Both *pkh1^ts^pkh2Δ* and *ypk1^ts^ypk2Δ* strains containing the plasmid pIRE1 were grown for 3 hours in the presence of DTT at both 25°C and 37°C. At both temperatures, the Ire1 protein levels were significantly reduced in the *pkh1^ts^pkh2Δ* strain (**Fig 3C** and **Supplemental Fig S7**). In the *ypk1^ts^ypk2Δ* cells, the Ire1 protein levels at 25°C were comparable to that of WT, while they were significantly reduced at 37°C (**Fig 3C** and **Supplemental Fig S7**). Similarly, the *pkc1^ts^* strain containing the plasmid pIRE1 was also grown for 3 hours with DTT at both 25°C and 38°C. At both temperatures, Ire1 levels were significantly reduced compared to WT cell (**Fig 3D** and **Supplemental Fig S7**). Collectively, these findings suggest that kinases Pkh1, Ypk1 and Pkc1 function in part within a common pathway and are required for maintaining the Ire1 protein abundance.

### 4. PKC1 and IRE1 are novel dosage suppressors of the *ypk1^ts^* strain

The tyrosine-to-cysteine mutation at residue 536 (Y536C) is responsible for the ts-phenotype of the Ypk1 kinase (34,35). Y536 is a conserved residue (corresponding Y288 in SGK1) and located at the C-lobe of kinase domain, a region likely involved in substrate or effector binding (**Supplemental Fig S8**). These observations prompted us to hypothesize that the Y536C substitution disrupts the binding of Ypk1 with one or more of its substrates or regulatory effectors, and that a high-dose of the proposed partners might rescue the ts-phenotype. Previous studies identified 8 dosage suppressors of the *ypk1^ts^ ypk2Δ* strain (39), including genes encoding the cell wall enzyme glucanase Exg1, the vacuolar protein sorter Sea4, and the ER-resident chaperone Hlj1 (**Supplemental Fig S9A)**. However, cells lacking either of those proteins grew on the medium containing tunicamycin (data not shown), suggesting that they had a minor role in ER stress response.

We also examined if these suppressors contributed to ER stress response when over-expressed in *ypk1^ts^ ypk2Δ* strain. The selected plasmids (**Supplemental Fig S9)** from the yeast genome tiling library containing the identified suppressor genes were introduced into the *ypk1^ts^ ypk2Δ* strain. The plasmid containing the *EXG1* and its neighboring genes restored growth of the *ypk1^ts^ ypk2Δ* strain at 37°C, but not in the presence of tunicamycin (**Supplemental Fig S9)**. These results indicated that increased expression of glucanase Exg1 can compensate for the reduce activity of Ypk1. However, the suppressive mechanism remains unclear.

To identify Ypk1-effectors involved in ER stress response, we conducted a dosage suppressor genetic screen under an ER stress condition. The tiling library of yeast genomic DNA was introduced into the *ypk1^ts^ ypk2Δ* strain. Transformants capable of growing at 37°C in the presence of tunicamycin were selected. Finally, the screen identified 5 plasmids (P1-P5), each of which restored the growth of *ypk1^ts^ ypk2Δ* strain at 37°C in the presence of tunicamycin (**Supplemental Fig S10)**. Further analysis of these plasmids showed that they contained coding sequence of the protein kinase Ypk1, Ypk2, or Pkc1, the GTPase Rho2, or the RNase Ire1(**Supplemental Fig S10)**.

To further characterize these suppressors, we individually over-expressed Ypk1, Pkc1, Ire1, Pkh1 (the upstream kinase of Ypk1) and Slt2 (the downstream effector of Pkc1) into the *ypk1^ts^ypk2Δ* strain using 2µ-plasmids: pGAL1-YPK1, pGAL1-PKC1, pIRE1, pGAL1-PKH1, and pGAL1-SLT2, respectively. Expressions of Ypk1, Pkc1, Pkh1 and Slt2 were driven by the *GAL1* promoter, whereas Ire1 were expressed under the control of its native promoter (**Supplemental Fig S11)**. As expected, the *ypk1^ts^ypk2Δ* strain expressing Ypk1 from the pGAL1-YPK1 grew at both 25°C and 37°C, regardless of the presence and absence of tunicamycin (**Fig 4A**, left panel). The same strain carrying plasmid pGAL1-PKC1, pGAL1-SLT2, or pIRE1, but not pGAL1-PKH1, grew at 37°C in the presence of tunicamycin (**Fig 4A**, left panel). Although the strain carrying the pIRE1 exhibited slow growth (**Fig 4A**, left panel), these findings demonstrate that Pkc1, Slt2 and Ire1 can compensate for the functional defect associated with the *ypk1^ts^* mutation. These findings further suggest that Ypk1, Pkc1 and Slt2 function within a common signaling pathway that promotes cellular adaptation to ER stress.

**Fig 4:**
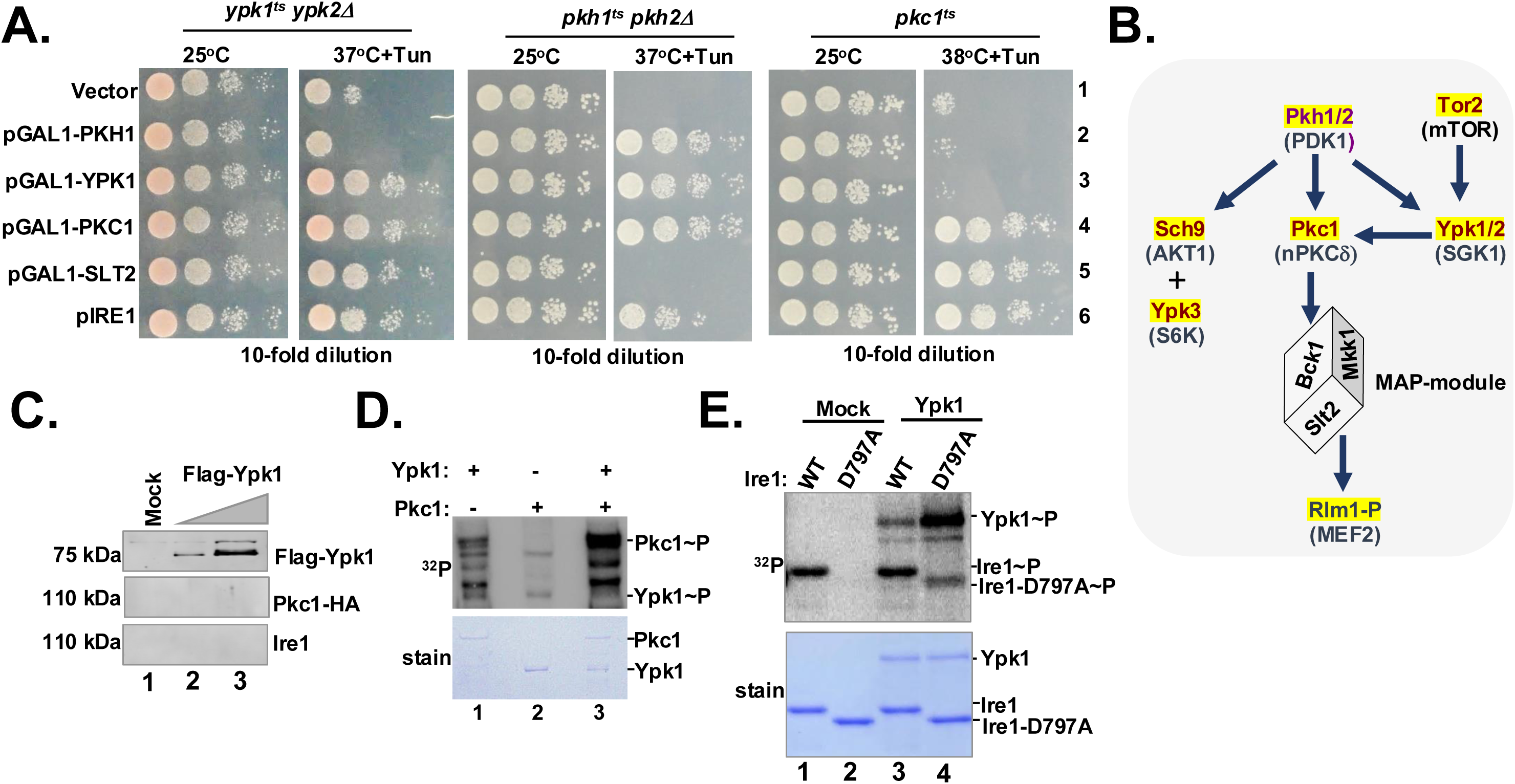
Over-expression of Pkc1 and Ire1 suppresses the temperature-sensitive phenotype of *pkh1^ts^* and *ypk1^ts^* strains. **(A)** The indicated yeast strains were transformed with the indicated plasmids expressing the indicated Pkh1, Ypk1, Pkc1, Slt2 or Ire1. Transformants were then tested for growth at permissive (25°C) and non-permissive (37°C or 38°C) temperatures in the presence and absence of tunicamycin (Tun). **(B)** The schematic of signaling pathways regulated by Pkh1, Ypk1 and Pkc1. **(C)** Flag-Ypk1 was co-expressed with either HA-tagged Pkc1 (Pkc1-HA) or Ire1 in yeast cells. Flag-Ypk1 was then immunoprecipitated by Flag-agarose and the associated proteins were detected by Western blot analysis, using anti-HA and anti-Ire1 antibodies. **(D)** & (**E**). Recombinant Ypk1, Pkc1 or Ire1 (WT and D797A) proteins were mixed in a kinase buffer containing the radioactive ^32^P-ATP for 15 minutes. The reaction was stopped by adding 2X SDS dye. The reaction products were separated in an SDS-PAGE, strained dried, and autoradiographed (^32^P).

We also introduced the above plasmids (i.e., pGAL1-PKH1, pGAL1-YPK1, pGAL1-PKC1, pGAL1-SLT2, and pIRE1) into *pkh1^ts^pkh2Δ* and *pkc1^ts^* strains. As expected, the plasmid expressing WT allele (pGAL1-PKH1 or pGAL1-PKC1) was able to support growth of the respective ts-strain at non-permissive temperatures, regardless of the presence and absence of tunicamycin (**Fig 4A**). Notably, over-expression of Ypk1 or Pkc1 from pGAL1-YPK1 and pGAL1-PKC1, respectively, supported the *pkh1^ts^pkh2Δ* strain to grow at 37°C in the presence of tunicamycin (**Fig 4A**, rows 3 and 4). These results suggest that increased Ypk1 and Pkc1 abundance can compensate for the reduced Pkh1 activity, consistent with Ypk1 and Pkc1 function downstream of Pkh1. In contrast, pGAL1-SLT2 failed to support the growth of *pkh1^ts^pkh2Δ* strain (**Fig 4A**, row 5), whereas pIRE1 conferred moderate support to grow at 37°C in the presence of tunicamycin (**Fig 4A**, row 6). These results suggest that the role of Slt2 is not limited to activating the Ire1 pathway.

The plasmids pGAL1-SLT2 or pIRE1 supported the *pkc1^ts^* strain to grow at 38°C in the presence of tunicamycin (**Fig 4A**, rows 5 and 6), suggesting that increased Slt2 or Ire1 abundance can compensate for the reduced Pkc1 activity, consistent with both Slt2 and Ire1 function downstream of Pkc1. Together, these results also suggest that a functional link exists between the Pkh1/2, Ypk1/2, Pkc1 and Ire1.

Previous studies have shown that Pkh1 directly phosphorylates and activates both Ypk1(39) and Pkc1(53). Additionally, Ypk1 is regulated by Tor2 complex(47,48), whereas Pkc1 acts upstream of Slt2(54). Our observations that both Pkc1 and Slt2 are dosage suppressors of *ypk1^ts^* allele, suggesting that Pkc1 and Slt2 likely function downstream of Ypk1. Collectively, these findings support a signaling hierarchy in which Pkh1 acts upstream of both Ypk1 and Pkc1, with Ypk1 regulating Pkc1 and its downstream effectors while itself being activated by an upstream kinase Tor2 (**Fig 4B**).

Based on the proposed signaling hierarchy, we proposed that Ypk1/2 regulate the UPR, at least in part, by functional association with Pkc1. The proposed Ypk1-Pkc1 complex either directly phosphorylates and activates Ire1 or indirectly activates Ire1 through one or more effectors, leading to *HAC1* mRNA splicing and UPR activation. Alternatively, the Ypk1-Pkc1 complex could stimulate the MAP kinase signaling cascade, including Slt2, which in turn promotes Ire1 expression though the transcription factor Rlm1 **(Fig 4B**), as we recently demonstrated(31).

### 5. Ypk1 phosphorylates Pkc1, but not Ire1

To investigate the physical associations among Ypk1, Pkc1, and Ire1, Flag-tagged Ypk1 [Flag-Ypk1] was co-expressed with either HA-tagged Pkc1 (Pkc1-HA) or Ire1 in yeast cells. Flag-Ypk1 was immunoprecipitated by Flag-agarose, and associated proteins were detected by Western blot analysis, using anti-HA, anti-Ire1 and anti-Slt2 antibodies. As expected, Flag-Ypk1 was readily detected (**Fig 4C)**. However, little or no Pkc1-HA and Ire1 was detected in the pulldown (**Fig 4C).** These results suggest that Ypk1 does not form a stable complex with Pkc1 or Ire1. Alternatively, these interactions are weak/or transient, as is often characteristic of kinase-substrate recognition(55).

To determine whether Ypk1 phosphorylates Ire1 or Pkc1, we performed *in-vitro* kinase assays. Flag-tagged Ypk1 and HA-tagged Pkc1 were expressed in yeast under the control of a galactose-inducible promoter and partially purified by Flag and HA-agarose, respectively. To be noted that the expression levels of both kinases were extremely low, making large-scale protein purification to homogeneity challenging. Therefore, partially purified recombinant Flag-Ypk1 protein was incubated with Pkc1-(HA)_3_, His_6_-Ire1^cyto^ (cytosolic domain of Ire1) or His_6_-Ire1^cyto^-D797A (kinase-inactive) in a reaction buffer containing radioactive ATP (^32^P-y-ATP). The reaction products were then separated by SDS-PAGE and subjected to autoradiography to detect the ^32^P incorporation in the kinases and their potential substrates.

The basal level of ^32^P incorporation was observed in Pkc1-(HA)_3_ and Flag-Ypk1 proteins when incubated with ^32^P-y-ATP alone (**Fig 4D** and **Supplemental Fig 12**), indicating that Pkc1-(HA)_3_ was phosphorylated itself. ^32^P incorporation into Pkc1-(HA)_3_ protein was markedly increased when incubated with wild-type Flag-Ypk1 (**Fig 4D**, lane 3). Two low-molecular weight radioactive protein bands below the Pkc1 band were also observed, which may represent degradation products of Pkc1-(HA)_3_ (**Fig 4D**, lane 3). Collectively, these results indicate that Flag-Ypk1 phosphorylated Pkc1-(HA)_3_.

^32^P incorporation into the Ire1^cyto^ protein in the absence Ypk1 kinase (**Fig 4E**, lane 1, Ire1^cyto^-P) indicated that Ire1^cyto^ protein was auto-phosphorylated. In contrast, no detectable ^32^P incorporation was observed in the kinase-dead Ire1^cyto^-D797A mutant protein (**Fig 4E**, lane 2, Ire1^cyto^-D797A-P), confirming its loss of kinase activity (56). When Ire1^cyto^-D797A was incubated with Ypk1, ^32^P incorporation was detected in both proteins (**Fig 4E**, lane 4, Ire1^cyto^-D797A-P). Interestingly, a basal level of ^32^P was incorporated in the re1^cyto^-D797A, while majority of the ^32^P was incorporated into Ypk1, reflecting its strong autophosphorylation activity (**Fig 4E**, lane 4, Ypk1-P).

In contrast, no detectable ^32^P incorporation was detected into Ire1 when it was incubated with the recombinant Pkh1 protein, indicating that Ire1 is not a direct substrate of Pkh1 (**Supplemental Fig S12**). Together, the basal phosphorylation of Ire1^cyto^-D797A protein and the relatively higher phosphorylation of Ypk1 indicate that Ire1 is unlikely to be a direct substrate of the kinase Ypk1. The low level of phosphorylation detected in Ire1^cyto^-D797A may instead results from an unidentified kinase associated with the Ypk1 preparation. Future work will be directed towards identifying this kinase and its potential role in Ire1 regulation.

### 6. Ire1-independent expression of Hac1 bypasses the need for Pkh1, Ypk1 and Pkc1

We investigated whether the requirement for Pkh1, Ypk1 and Pkc1 could be bypassed by expressing Hac1 independently of its cytosolic splicing. A *HAC1* variant harboring a G771A mutation in the intronic region was utilized for this purpose (**Fig 5A**). The *HAC1-G771A* variant bypasses the requirement of Ire1-mediated cytosolic splicing and produces a 230-amino-acid Hac1 isoform (Hac1^u^) from the un-spliced mRNA, with translation starting at the AUG codon of exon1 until the stop codon UGA at nucleotide 690 within the adjacent intron (57). As expected, the *ire1Δ* strain carrying a *HAC1*-G771A variant (**Fig 5B**, row 2), but not HAC1-WT (**Fig 5B**, row 1), grew on the tunicamycin medium at 25°C. The Hac1^u^ protein was expressed from the *HAC1*-G771A variant in the *ire1Δ* strain, which was enhanced when grown in the presence of DTT (**Fig 5C**, compare lanes 3 and 4). These results further validate our previous results that Hac1^u^ protein was expressed from the un-spliced mRNA(57).

**Fig 5:**
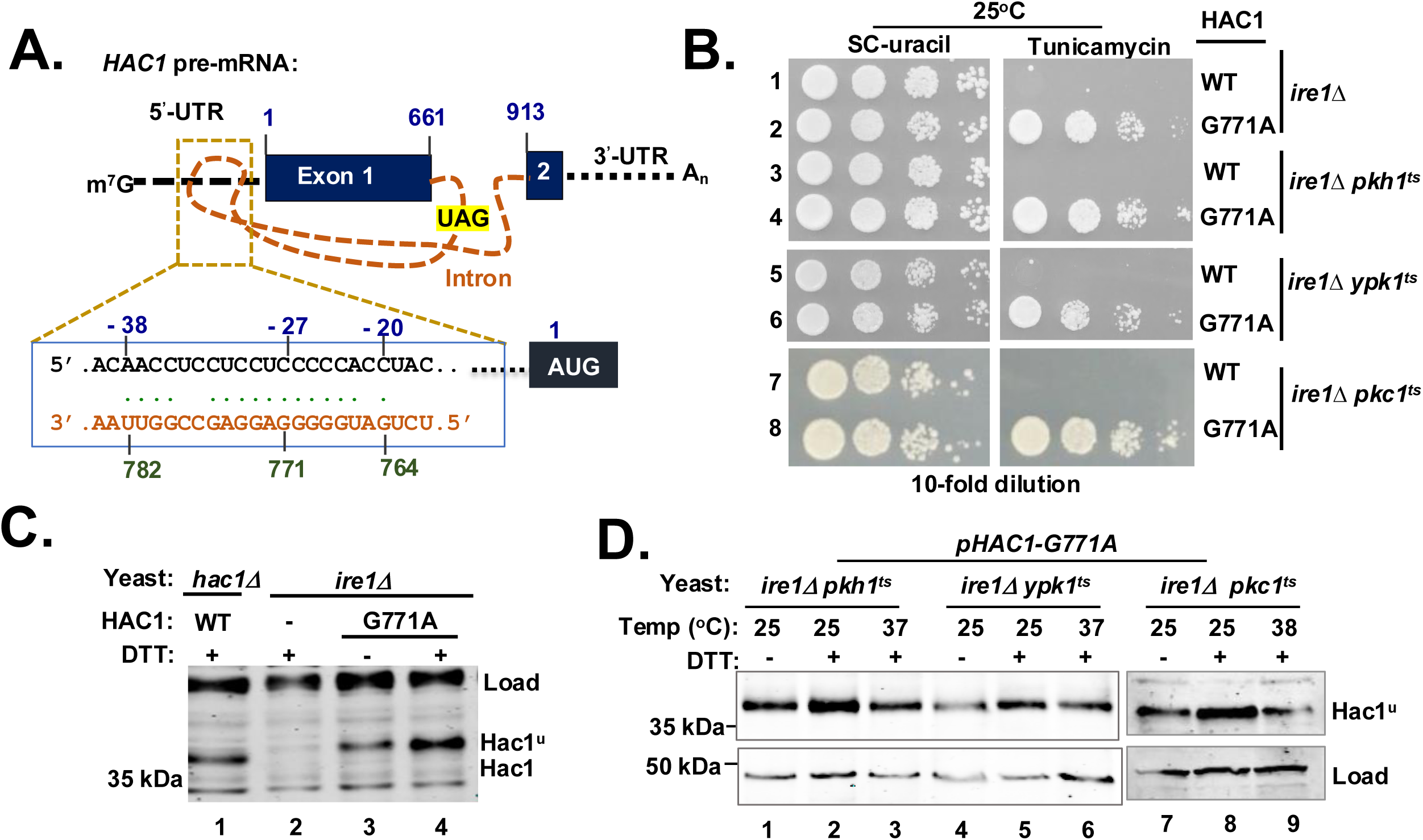
Bypassing the requirement of Pkh1, Ypk1 and Pkc1 for the Hac1-mediated ER stress response. **(A)** The schematic representation of *HAC1* pre-mRNA with 5’-7-methyl guanosine (m^7^G) cap and a poly-A tail (An). It includes solid rectangles representing the exons and an orange line representing the intron. The nucleotide compositions of the interaction between the 5’-UTR (5’-unstranslated region) and intron are indicated, with nucleotide numbers at the top. **(B)** The indicated yeast strains expressing type wild-type (WT) *HAC1* mRNA or *HAC1*-G771A mutant mRNA were serially diluted and grown on SC-uracil or the same medium containing tunicamycin (Tun) at 25°C. **(C)** WCEs were prepared from the indicated yeast strains in the presence and absence of DTT and subjected to Western blot analysis using antibody against the Hac1 protein. Hac1^u^ indicates Hac1 protein translated from the un-spliced mRNA. The non-specific bands are as a loading control. **(D)** The indicated yeast strains lacking Ire1 containing *HAC1-G771A* variant were grown in the presence or absence of DTT at 25°C, 37°C or 38°C. WCEs were prepared from and subjected to Western blot analysis using antibodies against Hac1 and Pgk1 proteins.

We disrupted the *ire1* gene in the genome of *pkh1^ts^pkh2Δ*, *ypk1^ts^ypk2Δ* and *pkc1^ts^* strains and created *ire1Δpkh1^ts^pkh2Δ*, *ire1Δypk1^ts^ypk2Δ*, *ire1Δpkc1^ts^* strains, respectively (**Supplemental Fig S13**). The resulting strains containing a WT-HAC1 allele did not grow on the tunicamycin medium at 25°C (**Fig 5B**, rows 3, 5, and 7). In contrast, they grew when containing a *HAC1*-G771A variant (**Fig 5B**). Consistent with this growth phenotype, Hac1^u^ protein expression was observed from the *HAC1*-G771A mRNA in *ire1Δpkh1^ts^pkh2Δ*, *ire1Δypk1^ts^ypk2Δ* and *ire1Δpkc1^ts^* strains when grown in the absence of DTT (**Fig 5D**). The Hac1^u^ protein expression was enhanced when cells were grown in the presence of DTT (**Fig 5D).** These results suggest that Pkh1, Ypk1 and Pkc1 had a minimum or marginal influence on the Hac1^u^ expression from the un-spliced *HAC1*-G771A mRNA. Together, our data suggest that Pkh1, Ypk1 and Pkc1 function to regulate *HAC1* mRNA splicing, but not Hac1 expression from the spliced mRNA.

### 7. Functional conservation between yeast Ypk1 and human SGK1

To assess the functional conservation between yeast Ypk1/2 and human SGK1, we introduced a plasmid expressing *SGK1* from a galactose-inducible promoter into both *ypk1Δ* and *ypk1^ts^ypk2Δ* strains. As reported earlier(58), the *ypk1Δ* strain expressing SGK1 fully complemented the Ypk1 function on the medium containing SFA (saturated fatty acid) and cerulenin (inhibitor of lipid biosynthesis) (**Supplemental Fig S2**). In contrast, SGK1 partially rescued the ts-phenotype of the *ypk1^ts^ypk2Δ* strain (**Fig 3**), despite robust SGK1 expression (**Fig 6**). The partial complementation may reflect differences in their domain architectures and/or distinct functional requirement under two different cell biological conditions. SGK1 consists of an N-terminal Phox-homology domain (PX), a central kinase domain and a C-terminal hydrophobic motif, whereas Ypk1/2 contains an N-terminal C2 domain, a central kinase domain, and C-terminal unstructured domain (**Fig 6** and **Supplemental Figure S2**).

**Fig 6:**
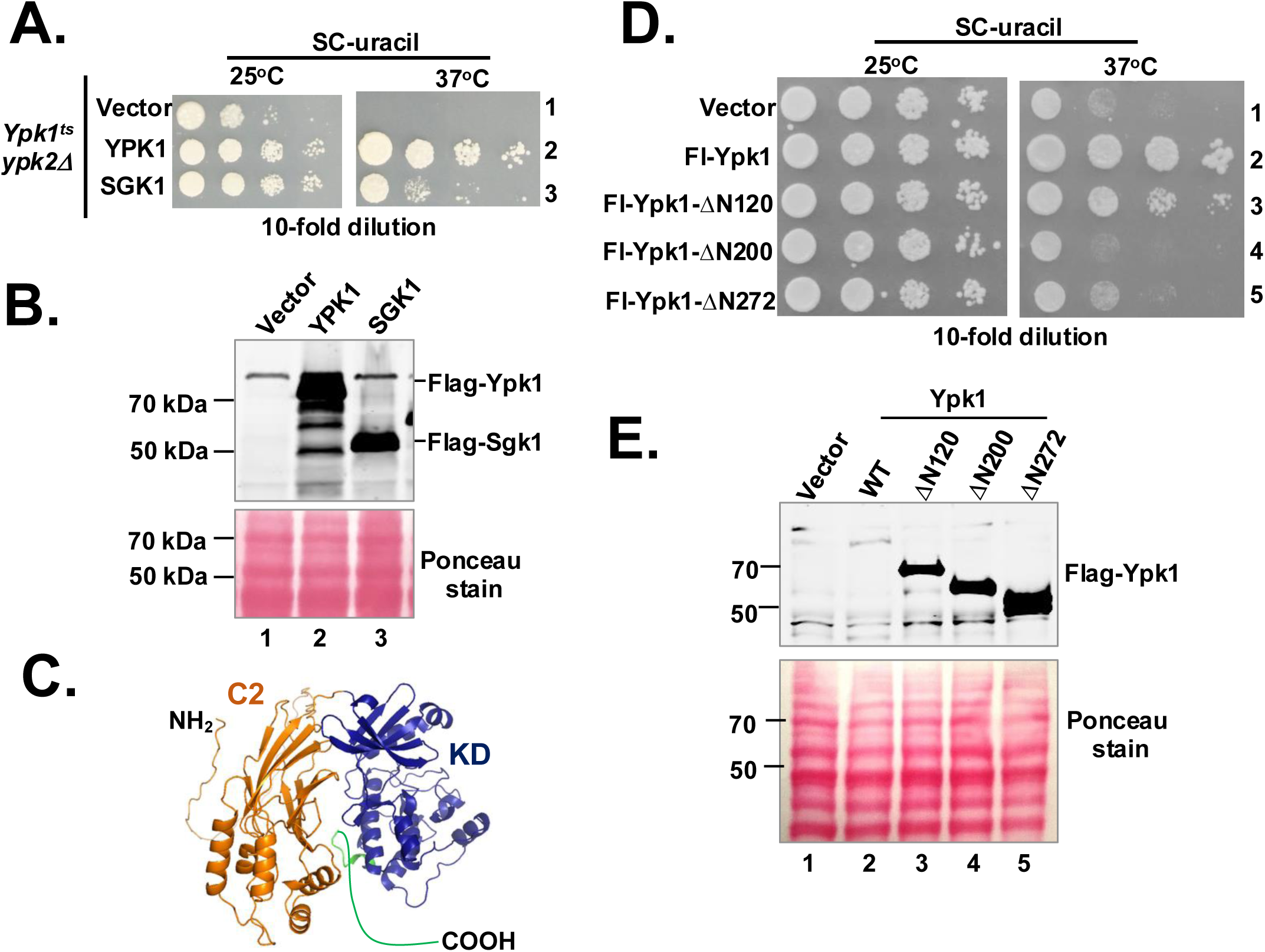
The C2-domain of Ypk1 is important for its function. **(A)** The *ypk1^ts^ypk2Δ* strain containing an empty vector or expressing YPk1 or SGK1 were serially diluted and grown at 25°C and 37°C for 48 hours. **(B)** WCEs were prepared from the indicated strains shown in (A) and subjected to Western blot analysis using anti-Flag antibody to detect Flag-Ypk1 (Fl-Ypk1) and Flag-SGK1 (Fl-SGK1) proteins. The Ponceau stained blot is shown as a loading control in the lower panel. **(C)** The AlphaFold predicted structure of the protein kinase Ypk1 was analyzed by PyMol software. Both C2 and kinase domains are indicated in orange and dark blue colors, respectively. **(D)** The *ypk1^ts^ypk2Δ* strains containing an empty vector or the same vector expressing the indicated Flag-tagged Ypk1 (Fl-Ypk1) and its indicated derivatives were serially diluted, spotted on the YEPD medium and grown at 25°C and 37°C for 72 hours. **(E)** WCEs were prepared from the indicated strains shown in (D) and subjected to Western blot analysis using anti-Flag antibodies to detect Flag-Ypk1 (Fl-Ypk1). The Ponceau stained blot is shown as a loading control in the lower panel.

The C2 domain belongs to a family of protein domains that mediate protein-protein interaction and target specific proteins to the cell membrane (59). At least 17 classes of C2 domains are known (60). According to the Alpha-fold prediction (61), the C2 domain in YPk1 (residues 116 to 340) is composed of nine β-strands sandwiched between five α-helices (**Fig 6C**). To determine its functional significance, we generated a series of deletion mutants. Specifically, 120, 200 and 272 amino acids were removed from the N-terminus of a Flag-tagged Ypk1 (Flag-Ypk1), generating Flag-Ypk1-ΔN120, Flag-Ypk1-ΔN200 and Flag-Ypk1-ΔN272 variants, respectively. These variants were then expressed into the *ypk1^ts^ypk2Δ* strain. Expression of the Ypk1 variants with an intact C2 domain (i.e., Flag-Ypk1 and Flag-Ypk1-ΔN120) fully complemented the Ypk1 function and restored the ts-phenotype, (**Fig 6D**, **Fig 6E** and **Supplemental Fig S14**). In contrast, variants lacking the C2 domain (i.e., Flag-Ypk1-ΔN200, and Flag-Ypk1-ΔN272) failed to restore the ts-phenotype (**Fig 6D**). These results suggest that the C2 domain is important for Ypk1 function, likely by promoting interaction with protein(s) required for its activity.

### 8. The IRE1 protein levels are reduced in SGK1-deficient human cells

To determine whether the Ire1 regulation by Ypk1 in yeast is conserved in mammalian cells, we examined human IRE1 expression in both HEK293FT and WA09 embryonic stem cells (ESC), following either SGK1 knockout (SGK1 KO) in HEK293FT (**Supplemental Figs S15 and S16**) or pharmacological inhibition of SGK1 using GSK650394 (5 µg/ml)(62). In both cell types, IRE1 protein levels were markedly reduced in the presence of 5 µg/ml of tunicamycin (**Figs. 7A** and **Supplemental Figs S17, S18 and S19**). Consistent with this decrease in IRE1 protein abundance, *XBP1* mRNA splicing (**Fig 7B** and **Supplemental Fig S20)** and XBP1 protein expression (**Fig 7A)** were significantly diminished in SGK1-KO cells as well as in HEK293FT and WA09 cells treated with both tunicamycin and GSK650394 **(Supplemental Figs S18 and S19**). These findings suggest that SGK1 contributes to UPR by modulating the abundance of IRE1 protein in human cells.

**Fig 7:**
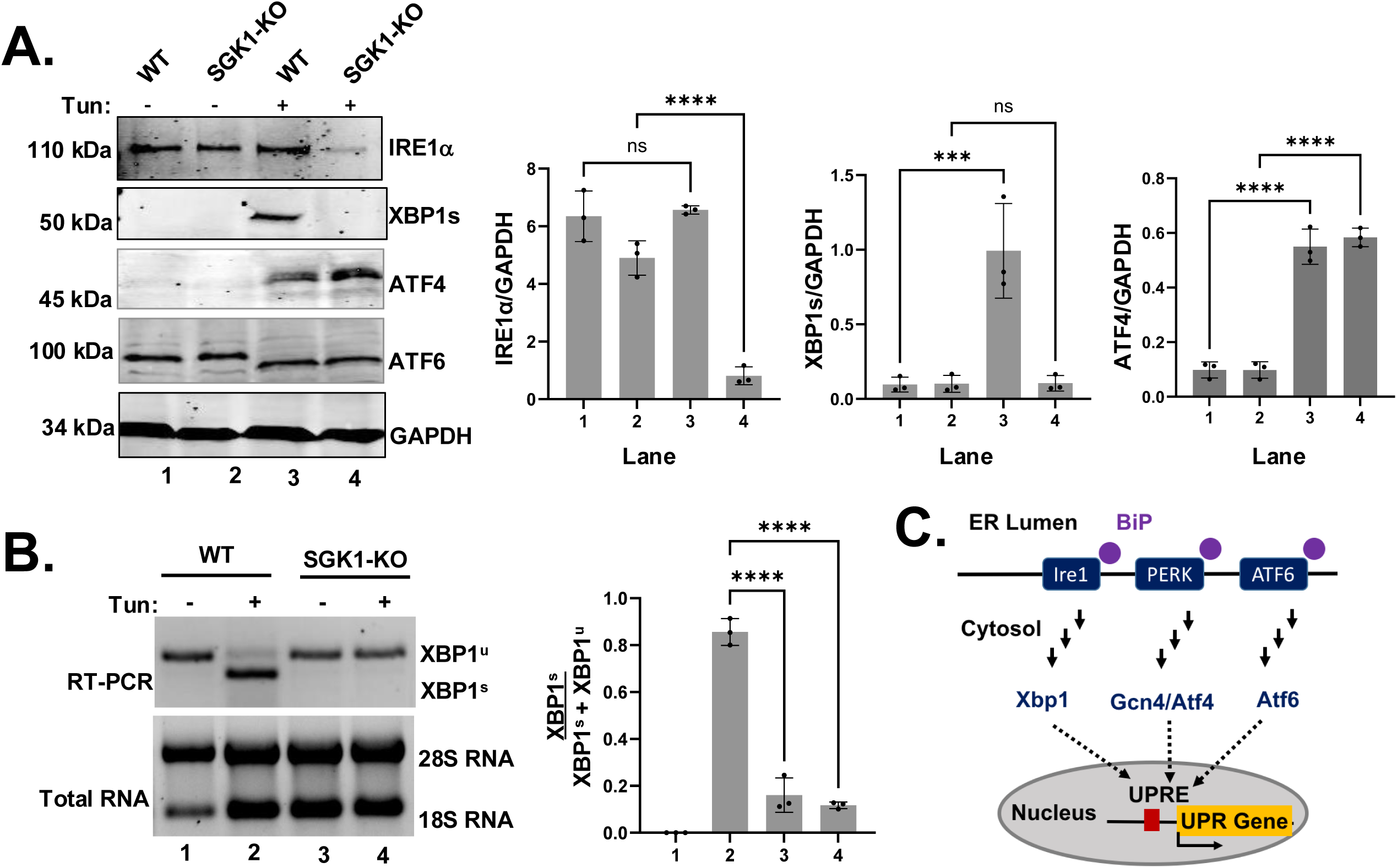
Reduce abundance of IRE1 protein and splicing of XBP1 mRNA in SGK1 knockout cells. **(A)** HEF293F cells lacking SGK1 was grown in presence (+) and absence (-) of tunicamycin (Tun). Whole cell extracts were prepared and subjected to Western blot analysis using indicated antibodies. Experiments were repeated at least three times. The intensities of protein bands were measured and shown in a bar diagram (****p-value<0.0001, paired t-test). **(B)** HEF293F cells lacking SGK1 was grown in presence (+) and absence (-) of tunicamycin (Tun). Total RNA was isolated and subjected to RT-PCR analysis to detect spliced (XBP1^s^) and un-spliced XBP1 (XBP1^u^) mRNA. Experiments were repeated at least three times. The intensities of RNA bands were measured and shown in a bar diagram (****p-value<0.0001, paired t-test). **(C)** Schematic representation of unfolded protein response-mediated by IRE1, PERK and ATF6. Unfolded protein is shown by sold circles

Because UPR is also regulated by two other major ER-stress sensors PERK and ATF6 (**Fig 7C**), we next examined the effects of SGK1 deficiency on those signaling branches. Specifically, we assessed the abundance of the ATF4 (a target of PERK) and ATF6 in the SGK1-deficient cells. Upon tunicamycin treatment, ATF4 protein level was increased in the SGK1 knockout (**Fig 7A)** or pharmacological inhibition (**Supplemental Figs S17, S18 and S19**). In addition, ATF6 exhibited faster mobility following tunicamycin treatment (**Fig 7A)**, consistent with its cleavage and activation(63). These data suggest that SGK1deficiency does not impair the PERK and ATF6 pathways. Together, these findings parallel our results in yeast and support a conserved mechanism in which SGK1 specifically regulates IRE1 abundance, while leaving the PERK and ATF6 branches of the UPR largely unaffected, thereby modulating the UPR activity.

## DISCUSSION

We show that yeast protein kinases Pkh1, Ypk1 and Pkc1 modulate the Ire1-mediated *HAC1* mRNA splicing and protein expression (**Fig 1 and Fig 2**), which is associated with the decreased Ire1 protein level (**Fig 3**). We also show that increased Ire1 abundance (**Fig 4**) or constitutive Hac1 expression (**Fig 5**) can compensate for the reduced activity of Pkh1, Ypk1 or Pkc1. Together, it appears that Pkh1/2, Ypk1/2, Pkc1 and Ire1 are functionally linked, which is supported by the previous report that Pkh1/2 phosphorylates Ypk1/2 (39) and our result that Ypk1 phosphorylates Pkc1 (**Fig 4**). Like yeast studies, the IRE1 protein levels in human cells were markedly reduced (**Fig 7**), when SGK1 function, the yeast ortholog of Ypk1 (**Fig 5** and **Fig S2**), was deleted or attenuated (**Fig 7**). This reduction in IRE1 abundance was accompanied by a significant decrease in *XBP1* mRNA splicing and XPB1 protein expression (**Fig 7**). Together, these results suggest that a conserved regulatory signaling Ypk1/1SGK1 pathway that promotes the *HAC1*/XBP1 mRNA splicing functions to regulate the Ire1/IRE1 protein abundance.

In metazoan cell, UPR is controlled by three major proteins (I) dual kinase/RNase Ire1(9–11,64), (II) kinase PERK(12) and (III) transcription factor ATF6(13). Ire1 initiates a signaling pathway by cleaving an intron from the translationally repressed mRNA *XBP1*(21,22,65) in human cells or *HAC1* in yeast cells(17,19,20,66). Cleaved exons of *HAC1* are joined by Rlg1 ligase(23) whereas cleaved exons of *XBP1* are joined by RTCB ligase(24), producing a matured *HAC1/XBP1* mRNA that translates the Hac1/XBP1 protein. PERK initiates a signaling pathway by phosphorylating the translation initiation factor 2α (eIF2α) leading to activation of the transcription factors Gcn4 in yeast cells and ATF4(67,68) in metazoan cells. ATF6 initiates a signaling pathway by relocating itself from the ER to the Golgi apparatus and releasing its transcriptional regulatory domain to the cytoplasm. The transcription factors XBP1, Gcn4 and ATF4 in turn increase the expressions of folding enzymes and chaperones (e.g., yeast Kar2 or human BiP(69–73)). In this study, we show that SGK1 specifically regulates IRE1 abundance (**Fig 7**), while leaving the PERK and ATF6 branches of the UPR largely unaffected.

The yeast *S. cerevisiae* genome encodes 129 protein kinases among which 19 are essential and 110 are nonessential (74). Among non-essential kinases 24 kinases have paralogs (**Supplemental Table 1**). Notably, deletion of both paralogs in Ypk1/2, Pkh1/2, Yck1/2, and Tor1/2 kinase pairs is lethal, suggesting that they collectively perform essential functions. Here, we examined 17 essential kinases and show that Cdc28, Pkc1, Rio2, Tor2, Pkh1, and Ypk1 play an important role in Ire1-mediated UPR (**Fig 1**). Our results demonstrate that Ypk1, Pkh1 and Pkc1 work in a shared, interconnected pathway and modulate the abundance of Ire1 protein (**Fig 6**). The roles of the remaining kinases (i.e., Cdc28, Rio2 and Tor2) in Ire1-mediated UPR remained to be determined.

The sequential steps in the Ire1-mediated UPR can be categorized as follows: (I) the abundance and activation of Ire1, (II) the colocalization of Ire1 and *HAC1*/*XBP1* mRNA, (III) the translation of spliced *HAC1/XBP1* mRNA, (IV) the translocation of Hac1/Xbp1 protein from cytosol to nuclear, and (V) the trans-activation of UPR genes. It is reasonable to assume that cells and organisms have evolved general and potentially gene-specific mechanisms to regulate each of these sequential steps of UPR. These molecular events are likely driven by dynamic changes in complex signaling networks involving both essential and non-essential kinases. Our understanding of these signaling networks, including their component, regulatory mechanisms, and their impact on the sequential steps of response the Ire1-Hac1/Xbp1 pathway to produce an additive response, remains incomplete.

The genetic screen and subsequent analyses revealed that over-expression of Pkc1 or Ire1 can compensate for the reduced function of Pkh1 and Ypk1 during ER stress (**Fig 4A**). These genetic observations provide strong evidence that Pkh1, Ypk1 and Pkc1 functionally contribute to ER stress response. Pkc1 is known to regulate the yeast MAPK module composed of MAP3K Bck1, MAP2K Mkk1/Mkk2, and MAPK Slt2(75,76). Slt2 has been shown to respond to ER stress by activating the Ire1 pathway(31), as well as by initiating a parallel signaling pathway(30). Previous studies have demonstrated that Pkh1/2 and Tor2 phosphorylate and activate Ypk1/2 (39) and our results show that Ypk1 phosphorylates Pkc1 (**Fig 4**). Together, these results support a model, in which that Pkh1 Ypk1 and Pkc1 function in a sequential signaling hierarchy to regulate Ire1 protein abundance. In this model (**Fig 8**), Pkh1 acts upstream of both Ypk1 and Pkc1, with Ypk1 regulating Pkc1 and its downstream effectors while itself being activated by an upstream kinase Tor2. The downstream effector of PKC is the MAPK module composed of MAP3K Bck1, MAP2K Mkk1/2 and MAPK Slt2 (77,78). The active Slt2 integrates with specific signaling molecules to reduce the cell wall stress (75,76) as well as ER stress through the transcription factor Rlm1(31) (**Fig 8**).

**Fig 8:**
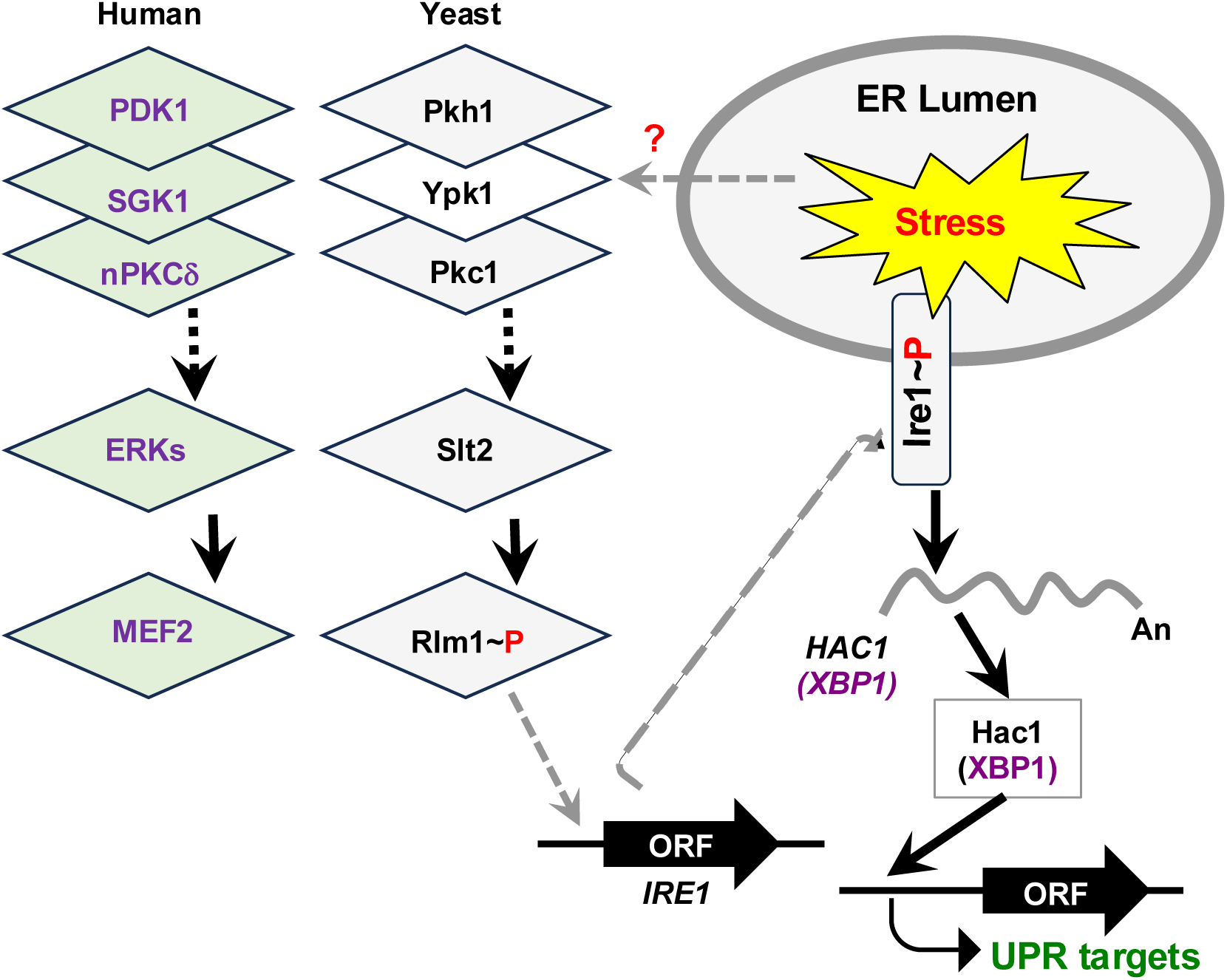
The conserved signaling hierarchy modulating Ire1 protein abundance. ER stress promotes Ire1 phosphorylation (Ire1∼P) and activation. Activated Ire1 initiates the adaptive UPR by mediating the cytosolic splicing of *HAC1/XBP1* mRNA (grey wavy line). Ire1 protein abundance is regulated by yeast a signaling hierarchy mediated by protein kinases Pkh1, Ypk1 and Pkc1, and Slt2, as well as the transcription factor Rlm1. The transmission of the unknow ER stress signal to the kinase complex is indicated by a red arrow. The corresponding human orthologs of yeast kinases and transcription factor are shown in parallel.

Human kinases PDK1, SGK1, mTOR, and nPKCδ are orthologs of yeast protein kinases Pkh1/2, Ypk1/2, Tor2 and Pkc1, respectively (**Fig 8**). PDK1 phosphorylates and activates at least 23 kinases, including SGK1(79). In addition to PDK1, mTOR also phosphorylates and activates SGK1, which then phosphorylates and activates numerous substrates, including NDRG1 (N-Myc downstream-regulated gene 1), FOXO3a (forkhead transcription factor 3a), NEDD4-2 (neuronal precursor cell expressed developmentally downregulated 4-2), thus regulating a broad range of biological events, such as cell growth and differentiation, apoptosis, and lipid homeostasis(80). In this study, we show that inhibition of SGK1 significantly reduces the cytosolic splicing of *XBP1* mRNA in three distinct human cell lines: HEK293FT, SGK1-KO in HEK293FT cells and WA09 embryonic stem cells. These results suggest that SGK1 plays a critical role in ER stress response and support the functional conservation of the Ypk1/SGK1-mediated regulation of the Ire1/IRE1 pathway from yeast to human cells. However, downstream signaling intermediates through which SGK1 regulates IRE1 abundance remain to be determined. Particularly, future studies are aimed at establishing if this regulation involves nPKCδ, NDRG1, FOXO3a, NEDD4-2, or other downstream effectors.

Here, our findings provide evidence that SGK1 regulates the IRE1 signaling pathway. Despite this functional conservation, Ypk1 and SGK1 exhibit important differences in their biological requirements. Ypk1 function is essential in yeast, whereas SGK1 function is not essential in human cells. The functional differences between yeast and human cells may, in part, reflect their distinct domain architectures (**Figs 3 and 6**). allowing each protein to associate with distinct binding partners, localize to different cellular compartment and undergo different mode of kinase activation. Consistent with this possibility, we found that the C2 domain is important for Ypk1 function (**Fig 6**), likely by promoting interaction with protein required for its activity. Thus, while Ypk1 and SGK1 have diverged in their domain organization and regulatory mechanisms, their ability to coordinate lipid homeostasis and stress-responsive signaling, including regulation of the Ire1/IRE1 pathway, appears to be evolutionarily conserved.

## EXPERIMENTAL PROCEDURES

### Yeast strains, growth, and gene disruption

Yeast *S. cerevisiae* strains were grown in the standard medium (1% yeast extract, 2% peptone, and 2% dextrose [YEPD]) or defined synthetic complete (SC, 0.17% yeast nitrogen base [YNB], 0.5% ammonium sulphate sulfate, 2% glucose or 2% galactose (wherever indicated) and all amino acids). The genomic DNA of the *ire1::hphMX* strain was used as a template to amplify the *hphMX* cassette using primers annealing ∼200-bases upstream and downstream of the *IRE1* open reading frame. The amplified PCR product was used to disrupt the *IRE1* gene of the *pkh1^ts^ pkh2Δ, ypk1^ts^ypk2Δ,* and *pkc1^ts^* strains. The list of yeast strains used in this study is shown in **Supplemental Table 2**.

### Dosage suppressor genetic screen of the ypk1^ts^ strain

A tiling library of yeast genomic DNA was introduced into the *ypk1^ts^ ypk2Δ* strain, and transformants capable of growing at 37°C were selected. 24 Plasmids were rescued from temperature-resistant colonies. 17 plasmids containing *YPK1* or *YPK2* gene were identified by PCR with gene-specific primers, while 7 other plasmids were characterized by end-sequence analysis. Three plasmids containing PKC1, RHO2 or IRE1 genes were identified (**Supplemental Fig S10**).

### Plasmids

Plasmids were made using standard molecular biological techniques. The list of plasmids used in this study is shown in **Supplemental Table 3**.

### Western blot analysis

Yeast cells were grown in YEPD or Synthetic Complete (SC) medium without appropriate nutrients until the OD_600_ value reached ∼0.5 to 0.6. DTT (5 mM) was added to the medium to induce ER stress, and cells were harvested after 2 or 3 hours (unless otherwise indicated). Whole-cell extracts (WCEs) were prepared by the TCA method as described previously (56). Proteins were then fractioned by SDS–PAGE, and Western blot analysis was performed using appropriate antibodies (**Supplemental Table 4**). All experiments were repeated at least two times.

### RNA analysis and reverse transcriptase (RT)-PCR of HAC1 mRNA

Yeast cells were grown in YEPD until they reached an OD_600_ value between 0.5 and 0.6. The ER stressor DTT (5 mM) was added to the medium and cells were grown further for another 2 hours. Cells were harvested, and RNA was isolated using the RNeasy mini kit (Qiagen). Purified RNA was quantified using a Nanodrop spectrophotometer (ND-1000, Thermo Scientific) and used to synthesize the first strand cDNA by a Superscript^TM^-III reverse transcriptase (Invitrogen 18080-093) and a reverse primer (5ʹ-CCCACCAACAGCGATAATAACGAG-3ʹ) that corresponded to nucleotides +1002 to 1025. To assay *HAC1* mRNA splicing, the synthetic cDNA was then PCR-amplified using a forward primer (5ʹ-CGCAATCGAACTTGGCTATCCCTA CC-3ʹ) that corresponded to nucleotides +35 to +60 and a reverse primer (5ʹ-CCCACCA ACAGCGATAATAACGAG-3ʹ) that corresponded to nucleotides +1002 to +1025. The PCR-amplified products were then run on a 1.5% agarose gel to separate spliced (HAC1^s^) and un-spliced (HAC1^u^) forms of *HAC1* mRNA. Quantities of HAC1^s^ and HAC1^u^ were measured using ImageJ. Experiments were repeated at least two times.

### Partial purification of proteins and in vitro protein kinase assay

Yeast cells (BJ2168) expressing the Flag-tagged Ypk1 protein or Pkc1-(HA)3 were grown in the presence of 10% galactose overnight. Cells were harvested, resuspended, and broken in a breaking buffer (20mM Tris-HCl pH 7.5, 500 mM NaCl, 0.1% Triton-X100, 1 EDTA-free protease tablet per 10 ml, 4µg/ml Leupeptin and 1 µM PMSF). The clear cell whole-cell lysate was prepared by centrifugation. The whole-cell lysate was incubated with Flag-agarose beads, and the Flag-Ypk1 protein was eluted by Flag-peptide (SIGMA, USA) in an elution buffer (20mM Tris-HCL pH 7.5, 50 mM NaCl, 10% glycerol and 10 mM DTT). The recombinant Pkc1-(HA)_3_ was purified by (HA)-agarose beads and HA-peptide (Thermo Scientific, USA).

The *E. coli* BL21(DE3) pLysE cells, bearing the plasmid D412 (expressing the His_6_-Ire1^cyto^) or D499 (expressing His_6_-Ire1^cyto-^D797A protein) were grown in the presence of 1 mM IPTG for overnight and His-tagged proteins were purified by standard protocol using Ni-agarose.

The in vitro kinase assay was performed with the partially purified Flag-Ypk1, Pkc1-(HA)3 and His_6_-Ire1 proteins in a reaction buffer (20 mM Tris-HCl, 50 mM KCl, 25 mM MgCl_2_, 1 µM PMSF, one protease tablet per 10 ml buffer and 1 µCi of ATP-[γ^32^P]. The reaction mixture was then separated by an SDS-PAGE. The gel was then stained, dried, and autoradiographed.

### Immunoprecipitation of Flag-Ypk1

The yeast strain (BJ2168) was transformed with two plasmids expressing Flag-tagged Ypk1 and HA-tagged Pkc1 (Pkc1-HA) or Ire1. Transformants were grown in the SGal (SC-uracil-leucin with 10% galactose) medium and whole cell extracts were prepared in buffer A (20mM Tris-HCl pH 7.5, 500 mM NaCl, 0.1% Triton-X100, 1 EDTA-free protease tablet per 10 ml, 4µg/ml Leupeptin and 1 µM PMSF). Flag-Ypk1 was immunoprecipitated by Flag-agarose, and associated proteins were detected by Western blot analysis with an appropriate antibody.

### Human cell culturing, SGK1 knockout and whole cell extract (WCE) preparation

HEK293FT (Invitrogen R70007) cells were cultured in a high glucose Dulbecco’s modified Eagle’s medium (DMEM) (Thermo Scientific) medium supplemented with 10% fetal bovine serum (Thermo Scientific), 100 U of penicillin G (Thermo Scientific), 100 µg/ml streptomycin (Thermo Scientific), and 6 mM L-glutamine (Thermo Scientific) as described before(31). Cells were treated with tunicamycin (5 µg/ml) for 4 hours and washed with 1X phosphate buffered saline (PBS) (Thermo Scientific). Cells were incubated in a lysis buffer (20 mM Tris-HCl (pH 8.0), 80 mM KCl, 1 mM EDTA, 0.5% NP-40, 1 mM DTT, one phosphatase inhibitor cocktail per 50 ml buffer) for 10 min and lysed by pipetting followed by vortexing for 20 min at 4°C. WCEs was prepared by centrifugation at 12,000 g for 20 min at 4°C and subjected to Western blot analysis using appropriate antibody (**Supplemental Table 5).**

Human WA09 embryonic stem cells (hESCs; Wicell Research Institute, Madison, Wisconsin, USA), licensed to Prof. Nadege N Gouignard^1^) with a normal karyotype were used as wildtype hESCs line. Cells were maintained on growth-factor reduced Geltrex-coated dishes (Gibco, #A1413302) in mTeSR™1 medium (Stem Cell Technologies, #85850) and cultured until they reached ∼80% confluence. The culture medium was replaced daily, cells were split every 5–7 days with accutase (Stem Cell Technologies, #07920)33 and replated at a 1:6 dilution onto the Geltrex-coated plates.

The sgRNAs targeting SGK1 were designed using the CRISPR Design Tool (crispr.mit.edu). To generate CRISPR/Cas9-mediated editing of SGK1, we constructed the plasmid D3004 by annealing two complementary oligonucleotides encoding the sgRNAs and cloning them into the BbsI-digested pSpCas9(BB)-2A-Puro (PX459) V2.0 vector (**Supplementary Fig S15**). HEK293FT cells were cultured in Dulbecco’s Modified Eagle’s Medium (DMEM) supplemented with 10% FBS and transfected with the plasmid D3004, using the Xtremegene transfection reagent(81). The SGK1 knockout cells were confirmed by T7 endonuclease assay.

### RNA isolation and reverse transcriptase (RT)-PCR of XBP1 mRNA

HEK293FT cells were cultured in high-glucose DMEM supplemented with 10% fetal bovine serum, 100 U/ml penicillin G, 100 µg/ml streptomycin, and 6 mM L-glutamine as described previously. Cells were treated with tunicamycin (5 µg/ml). Total RNA was isolated using QIAzol Lysis Reagent (QIAGEN, cat. no. 79306) Briefly, cells were lysed in QIAzol Lysis Reagent, followed by phase separation with chloroform. The aqueous phase containing RNA was recovered and RNA was precipitated with isopropanol. The RNA pellet was washed with 75% ethanol, air-dried, and dissolved in RNase-free water.

For first-strand cDNA synthesis, equal amounts of purified RNA were reverse-transcribed using SuperScript III reverse transcriptase and random hexamers according to the manufacturer’s instructions. The resulting cDNA was used as a template for PCR amplification using *XBP1*-specific forward primer (5′-CCTGGTTGCTGAAGAGGAGG-3′) and reverse primer (5′-CCATGGGGAGATGTT CTGGAGG-3′). PCR products were resolved on a 3% agarose gel to distinguish the spliced (*XBP1^s^*) and un-spliced (*XBP1^u^*) forms of *XBP1* mRNA. Gels were imaged using LiCOR Bio CCD Imager, and the relative abundance of *XBP1^s^* and *XBP1^u^* were quantified using ImageJ.

### Structural analyses

The coordinates of the predicted Ypk1 structure were retrieved from the AlphaFold web site (https://alphafold.ebi.ac.uk)(61) and analyzed using the free PyMol software (https://www.pymol.org/pymol)

### Data analysis and densitometry analysis

The densitometry analysis was performed using the NIH ImageJ gel analysis software(82). All bands at the correct molecular weight ± approximately 5 kDa were analyzed thrice as the signal for that target protein. The average of three measurements with SD (standard deviation) are shown in each plot. Comparisons are made against the same sample lysate. Each Western blot was repeated at least twice.

### Statistical Analysis

All quantitative experiments reported in this study were conducted using a minimum of three independent biological replicates (n = 3). For each biological replicate, yeast strains were independently cultured and treated under identical experimental conditions prior to sample collection.

All quantitative data were plotted and statistically analyzed using GraphPad Prism (version 10.1.1). Statistical parameters—including mean, standard deviation (SD), standard error of the mean (SEM), and *P* values—were calculated in GraphPad Prism. Statistical significance was assessed using paired Student’s *t* test, one-way ANOVA, or two-way ANOVA, as appropriate.

## DATA AVAILABILITY

The numerical data used for generating graphs are available in the Supplementary information, and the uncropped images can be found in the Supplementary Figure S22. All Additional data and resources, including plasmids, can be obtained from the corresponding author upon reasonable request.

## AUTHOR CONTRIBUTIONS

SC conceptualized, performed experiments, and wrote the paper; AC conceptualized, performed experiments, and wrote the paper; JKU conceptualized, performed experiments, and wrote the paper; NB performed experiments; NR conceptualized and guided protein purification; NG guided mammalian cell experiments; APTN guided the mammalian cell experiments; MD conceptualized, performed experiments, and wrote the paper. All authors edited the paper.

## ACKNOWLEDGMENTS

We would like to thank Professors Brenda Andrews (Department of Molecular Genetics, University of Toronto, Canada), Jeremy Thorner (UC Berkeley, California, USA), Michael N Hall (University of Basel, Switzerland), Robert C. Dickson (University of Kentucky College of Medicine, USA), and Suresh Subramani (UCSD, California, USA) for their generous help in providing the temperature-sensitive strains. We would like to thank Prof. Ted Power (UC Davis, California, USA) for providing us with some Ypk1 constructs. We would also like to thank William Gambon for help split the WA09 embryonic stem cells. This work was supported by grants to M.D. from the U.S. National Institutes of Health (R15 GM159328) and UWM graduate school (ARC grant).

## Supplemental Figures

**Supplemental Fig 1:**
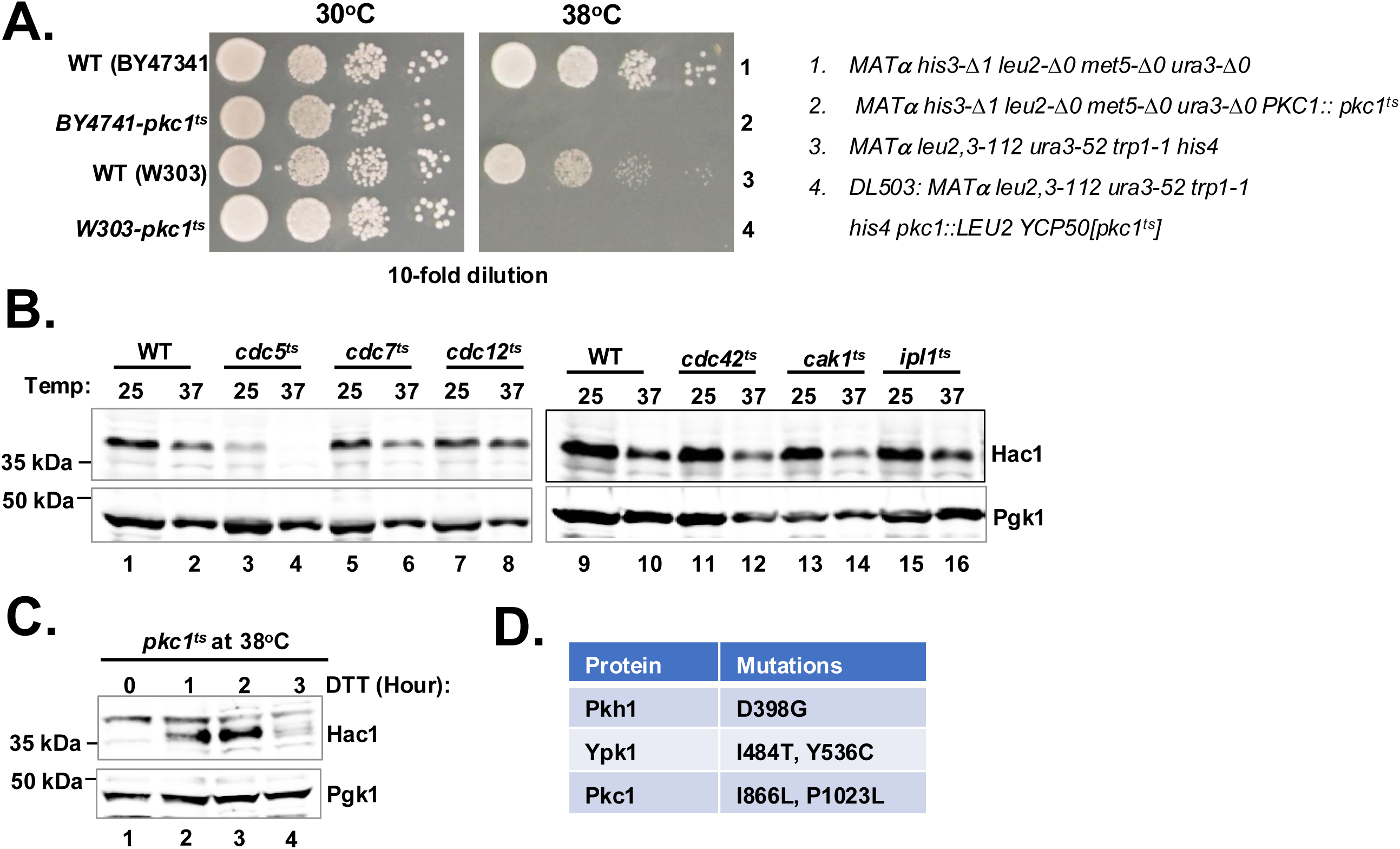
Analysis of Hac1 protein expression in temperature sensitive strain. (A) The indicated yeast *pkc1^ts^* strains were tested for growth on YEPD medium at 30°C and 38°C. The genotypes of yeast strain are illustrated. (B) Whole cell extracts were prepared from the indicated yeast strains and subjected to Western blot analysis using Hac1 and Pgk1 antibodies. (C) The *pkc1^ts^* strain was grown at 38°C in the presence of DTT for 1, 2 and 3 hours. Whole cell extracts were prepared and subjected to Western blot analysis using Hac1 and Pgk1 antibodies. (D) Table shows the Temperature-sensitive mutations in the indicated protein kinases

**Supplemental Fig 2:**
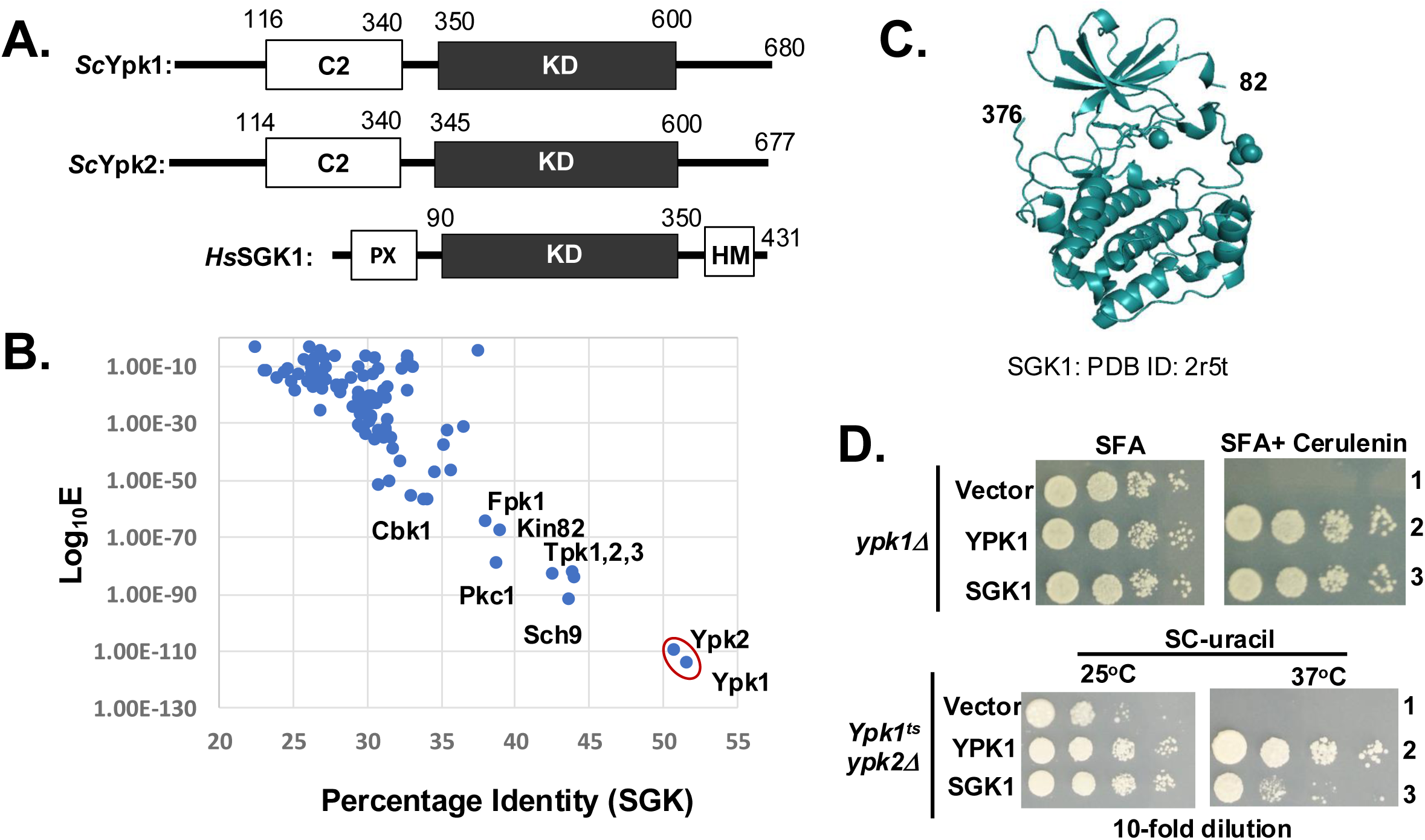
Analysis of Protein kinases Ypk1, Ypk2 and its human ortholog SGK1. **(A)** Schematic representations of *Saccharomyces cerevisiae* Ypk1 and Ypk2 (ScYpk1 and ScYpk2) and *Homo sapiens* SGK1 (HsSGK1). The C2 domain, kinase domain (KD) and hydrophobic motif (HM) are indicated The number indicates amino acid residues. (B) The protein sequence of human SGK1 was subjected to NCBI BLAST search against the *Saccharomyces cerevisiae* genome data base. The percentage identity and Log_10_E values are plotted. (C) The cartoon representation of SGK1 (PDB ID = 2R5T). (D) The indicated yeast strains expressing YPK or SGK1 were tested for growth on the indicated medium

**Supplemental Fig 3:**
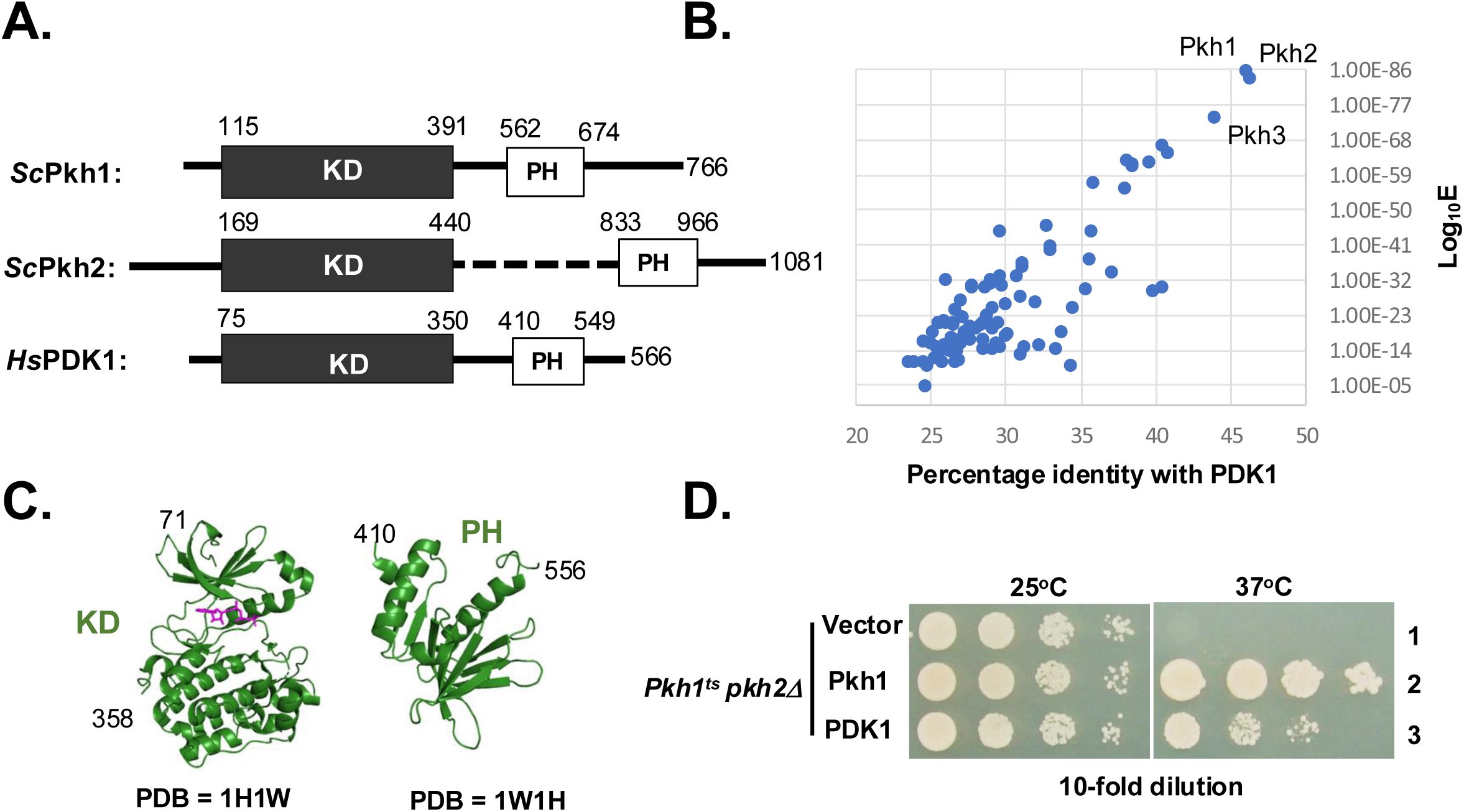
Protein kinases Pkh1, Pkh2 and its human ortholog PDK1. (A) Schematic representations of *Saccharomyces cerevisiae Pkh*1 and Pkh22 (ScPkh1 and ScPkh2) and *Homo sapiens PDK*1 (HsPDK1). The number indicates amino acid residues. (B) The protein sequence of human PDK1 was subjected to NCBI BLAST search against the *Saccharomyces cerevisiae* genome data base. The percentage identity and Log_10_E values are plotted. (C) The cartoon representation of the kinase and PH domains of PDK1 (PDB ID = 1H1W and 1w1H) (D) The indicated yeast strains expressing Pkh1 or PDK1 were tested for growth at 25°C and 37°C.

**Supplemental Fig 4:**
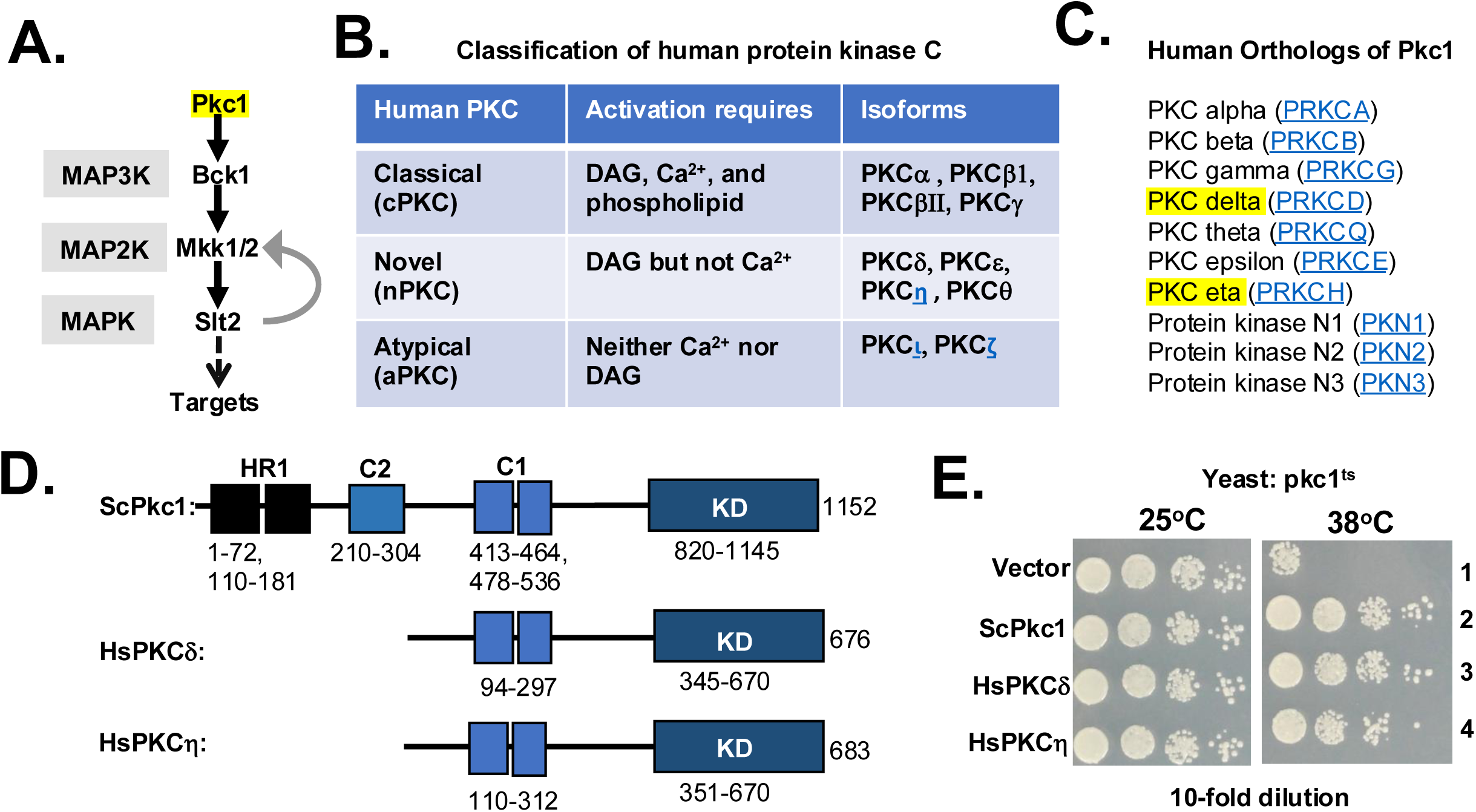
Protein kinase Pkc1 and its human orthologs PKCδ and PKCη. (A) Schematic of Slt2 MAPK signaling pathway. (B) Isoforms of protein kinase C (PKC) in human. (C) Human orthologs of yeast Pkc1. (D) Schematic representations of *Saccharomyces cerevisiae Pkc1 (*ScPkc1), Homo sapiens PKC-delta (HsPKCδ) and Homo sapiens PKC-eta (HsPKCη). The domains are shown in boxes. The number indicates amino acids. (E) The *pkc1^ts^* strains expressing the indicted human PKC alleles were tested for growth at 25°C and 38°C.

**Supplemental Fig 5:**
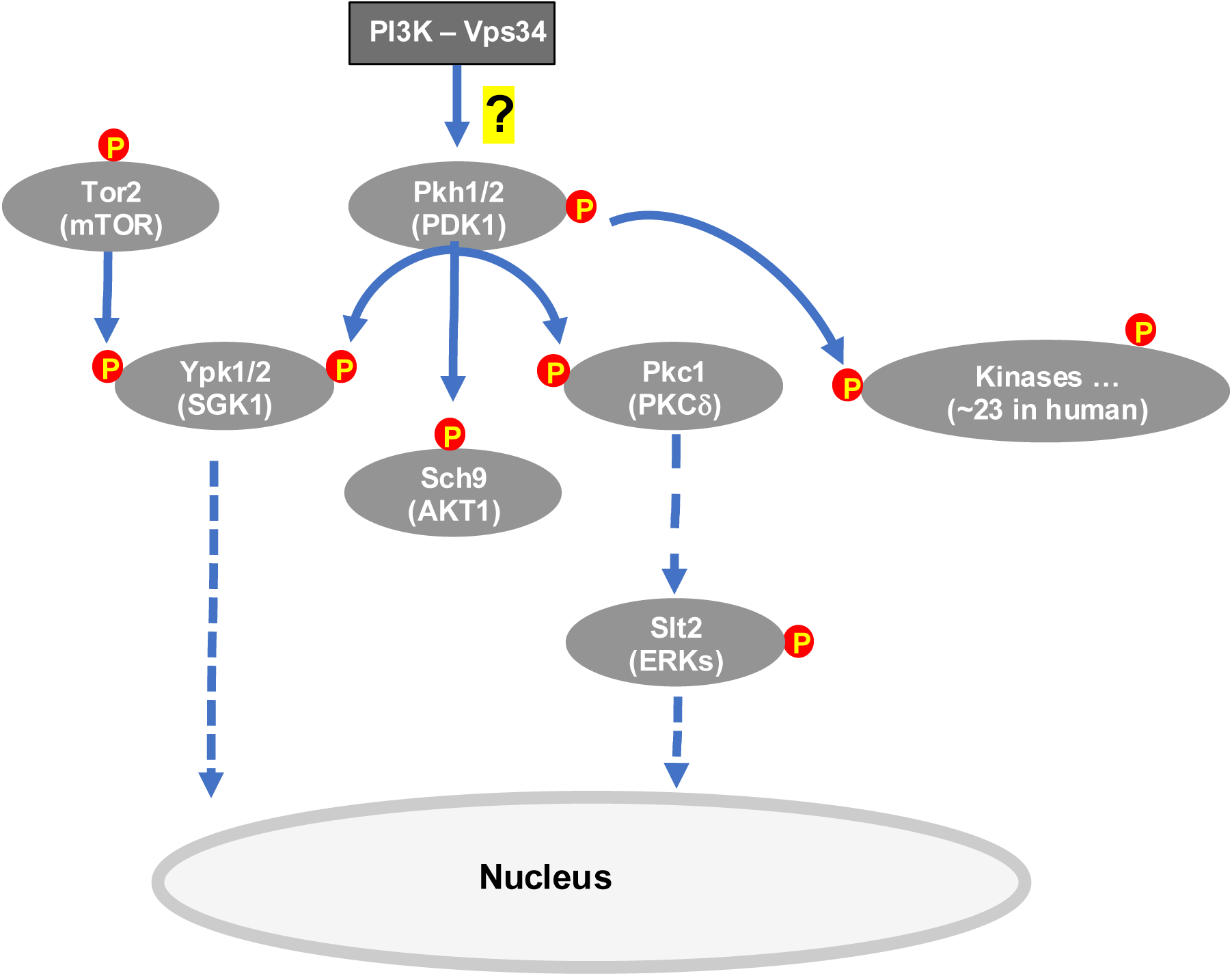
Protein kinases Ypk1, Pkh1 and Pck1 in the Signaling network. The model of Ypk1/2 and Slt2 protein kinase signaling pathway. The phosphorylation of the kinases are indicated by red circle. The human protein are bracketed.

**Supplemental Fig 6:**
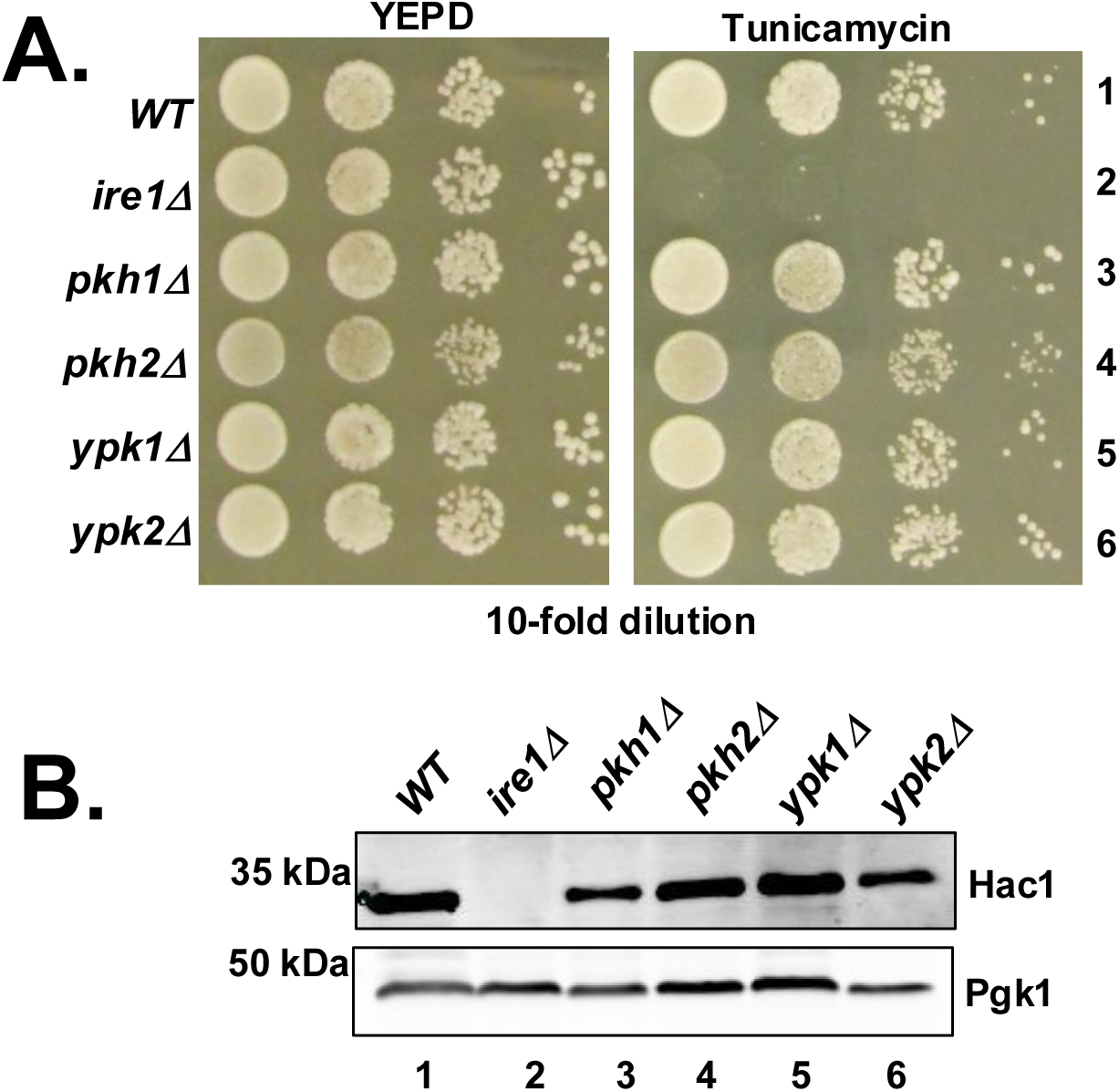
Analysis of Hac1 protein expression. (A) The indicated yeast *pkc1^ts^* strains were tested for growth on YEPD medium at 30°C (B) Whole cell extracts were prepared from the indicated strains in the presence of DTT and subjected to Western blot analysis by Hac1 and Pgk1 antibodies

**Supplemental Fig 7:**
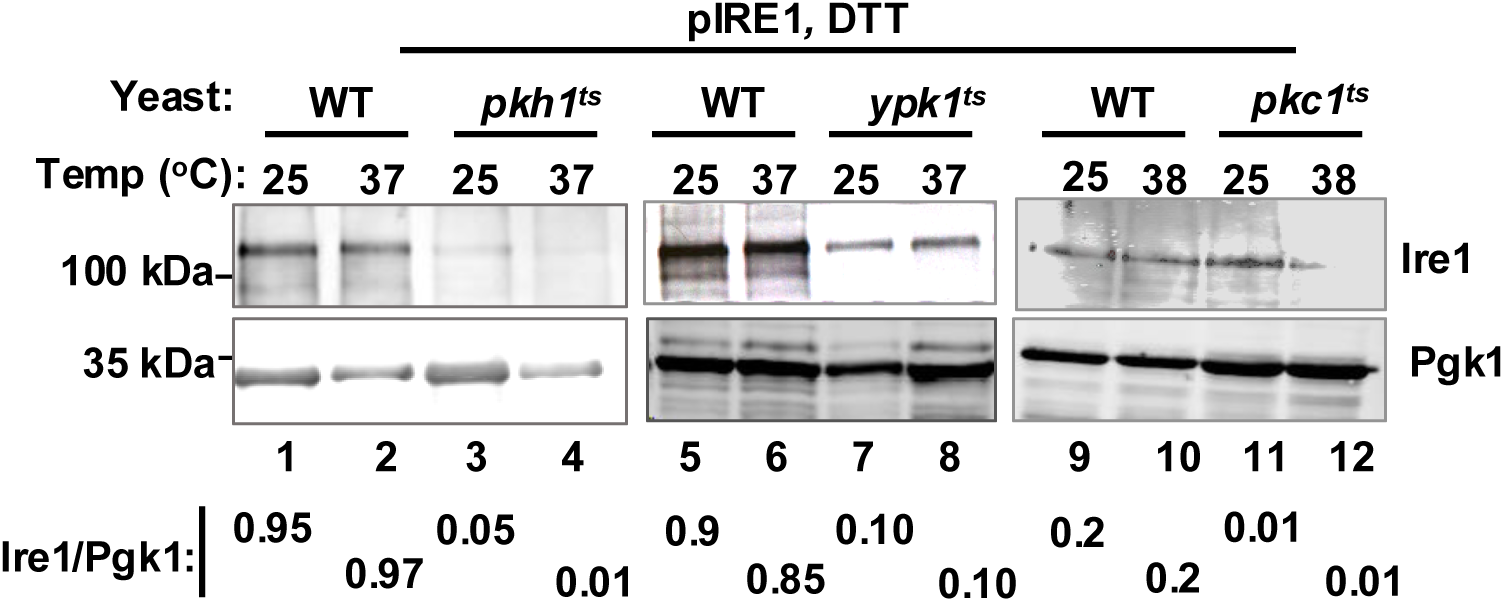
Analysis of Ire1 protein expression. The indicated yeast strains were transformed with a 2μ plasmid expressing Ire1 from its native promoter (pIRE1). Transformants were in the presence DTT at the indicated temperatures. Whole cell extracts were prepared and subjected to Western blot analysis by Ire1 and Pgk1 antibodies. The protein band intensities were measured. The ratios are indicated at the bottom.

**Supplemental Figure 8:**
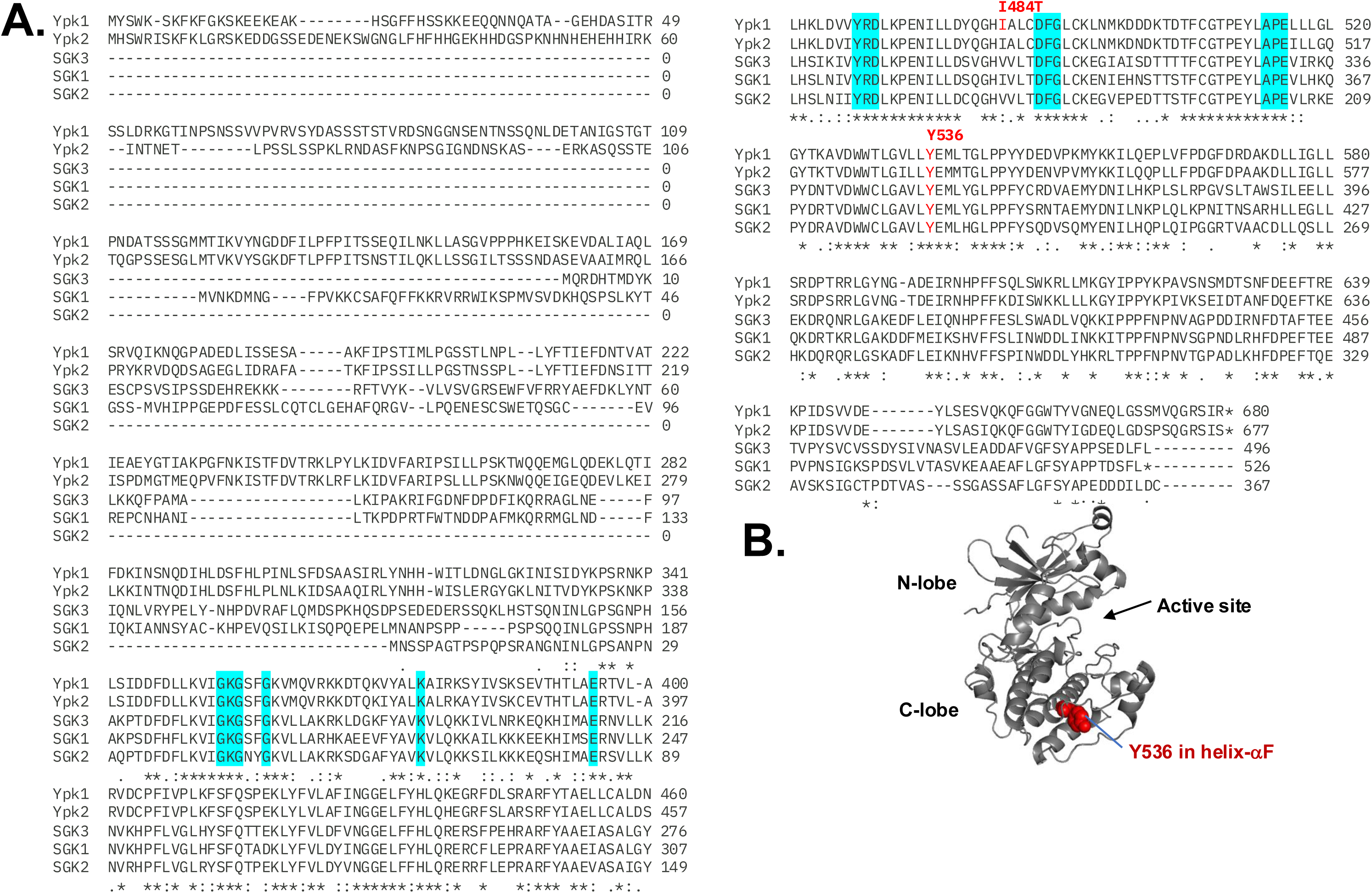
Protein sequence alignment of yeast Ypk1 and Ypk2 and their human orthologs SGK1, SGK2 and SGK3. **(A)** Multiple sequence alignment of protein kinases Ypk1/2 and its human orthologs SGK. Kinase conserved motifs are highlighted in blue. The mutations causing temperature-sensitive are indicated. **(B)** The Alpha-Fold predicted structure of the Ypk1 kinase domain.

**Supplemental Fig 9:**
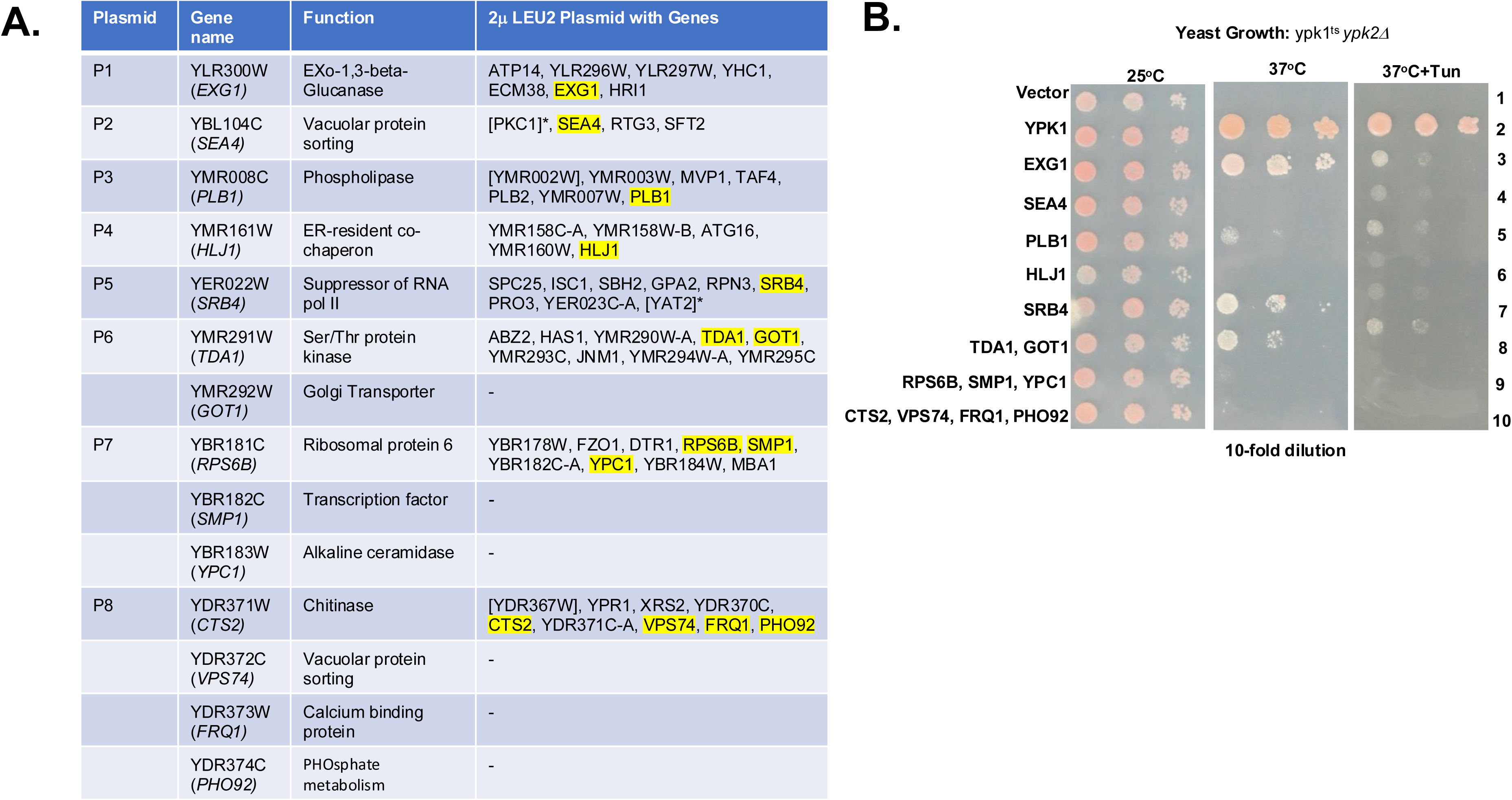
Published dosage suppressors of ypk1^ts^ strain. **(A)** The list of **p**lasmids (P1-P8) containing the genes corresponding to the reported high-copy suppressors of the *ypk1^ts^ ypk2Δ* strain. The plasmids also contain other indicated genes. (**B**) The *ypk1^ts^* strain containing the plasmids P1-P8 were tested for growth at the indicated temperatures in the presence or absence of tunicamycin (Tun).

**Supplemental Fig 10:**
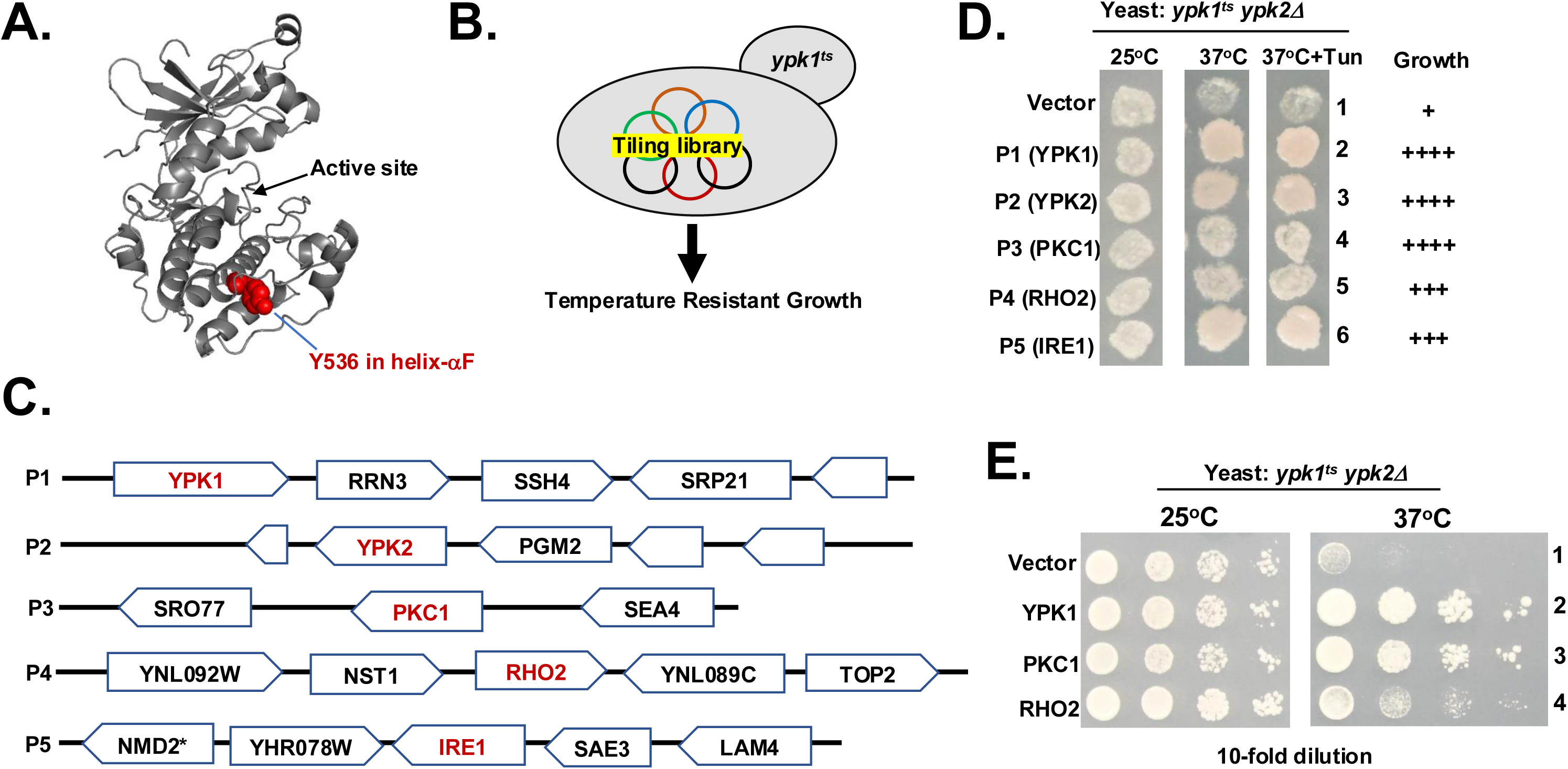
Genetic screen to identify suppressors of ypk1^ts^ strain. (A) The cartoon representation of the Ypk1 kinase domain (Alpha-fold predicted structure). Residue Y536 is shown in red. (B) The schematic of genetic screen, in which ypk1^ts^ strain was transformed with a high-copy genomic tiling library in a Leu2 plasmid. (C) Schematic representation of five plasmids (P1-P5) obtained from the genetic screen. Each plasmid harbors multiple genes which are shown in open arrows. (D) The *ypk1^ts^ ypk2Δ* strain containing the indicated plasmid were tested for growth with or without tunicamycin (Tun) at 25°C and 37°C. (E) The *ypk1^ts^ ypk2Δ* strain expressing Ypk1, PKC1 or Rho2 from a 2μ plasmid from a GAL1 promoter were serially diluted and tested for growth at 25°C and 37°C.

**Supplemental Fig 11:**
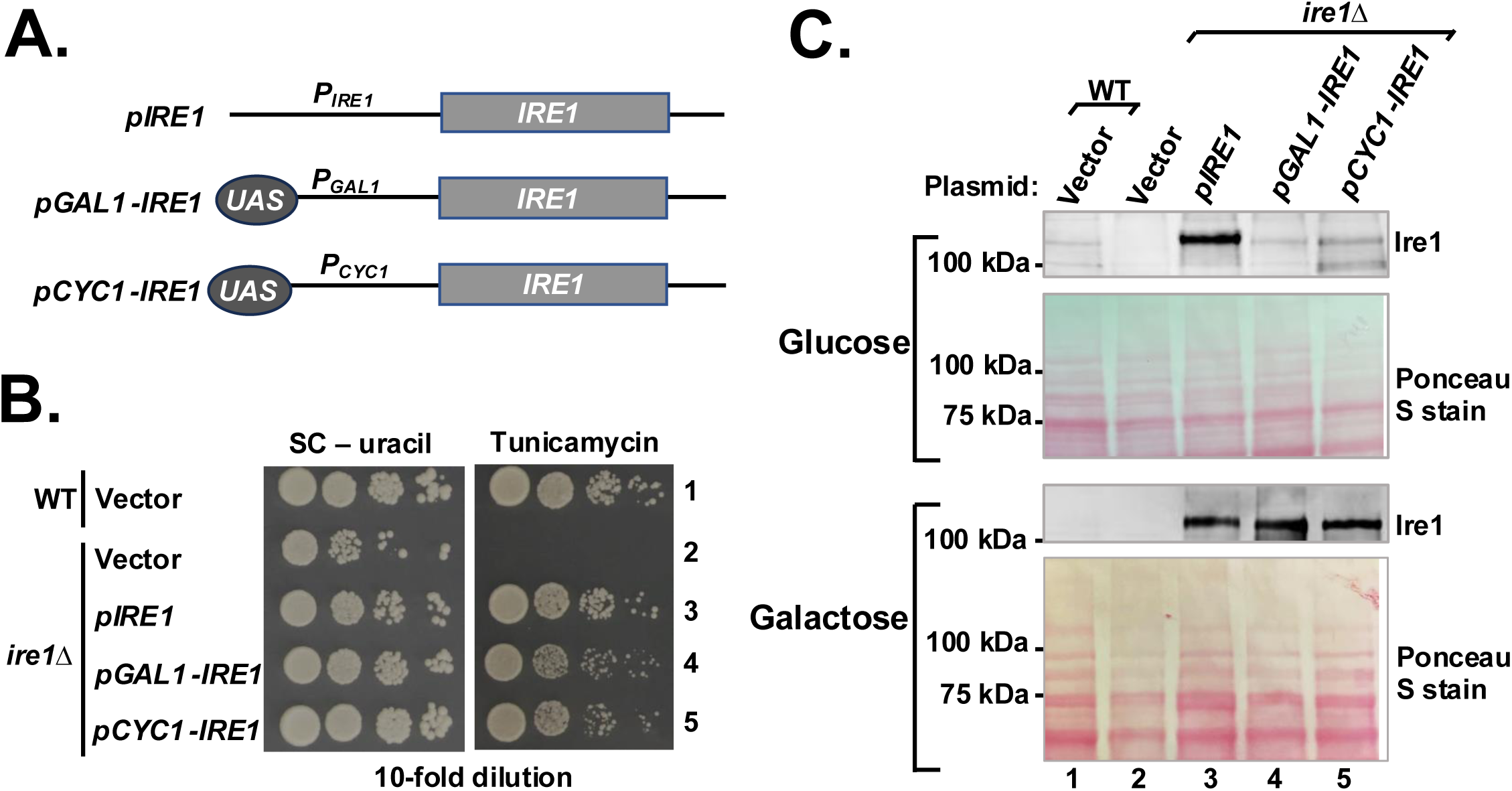
Ire1 expressed from the GAL1 or CYC1 promoter. (A) Schematic of IRE1 constructs. (B) WT and ire1Δ strains containing the vector plasmid or the same plasmid expressing IRE1 were tested for growth on synthetic complete (SC) medium and the same medium with tunicamycin. (C) Yeast strains as shown in (A) were grown in glucose and galactose medium. Whole cell extracts were prepared and subjected to Western blot analysis using Ire1 antibody.

**Supplemental Fig S12:**
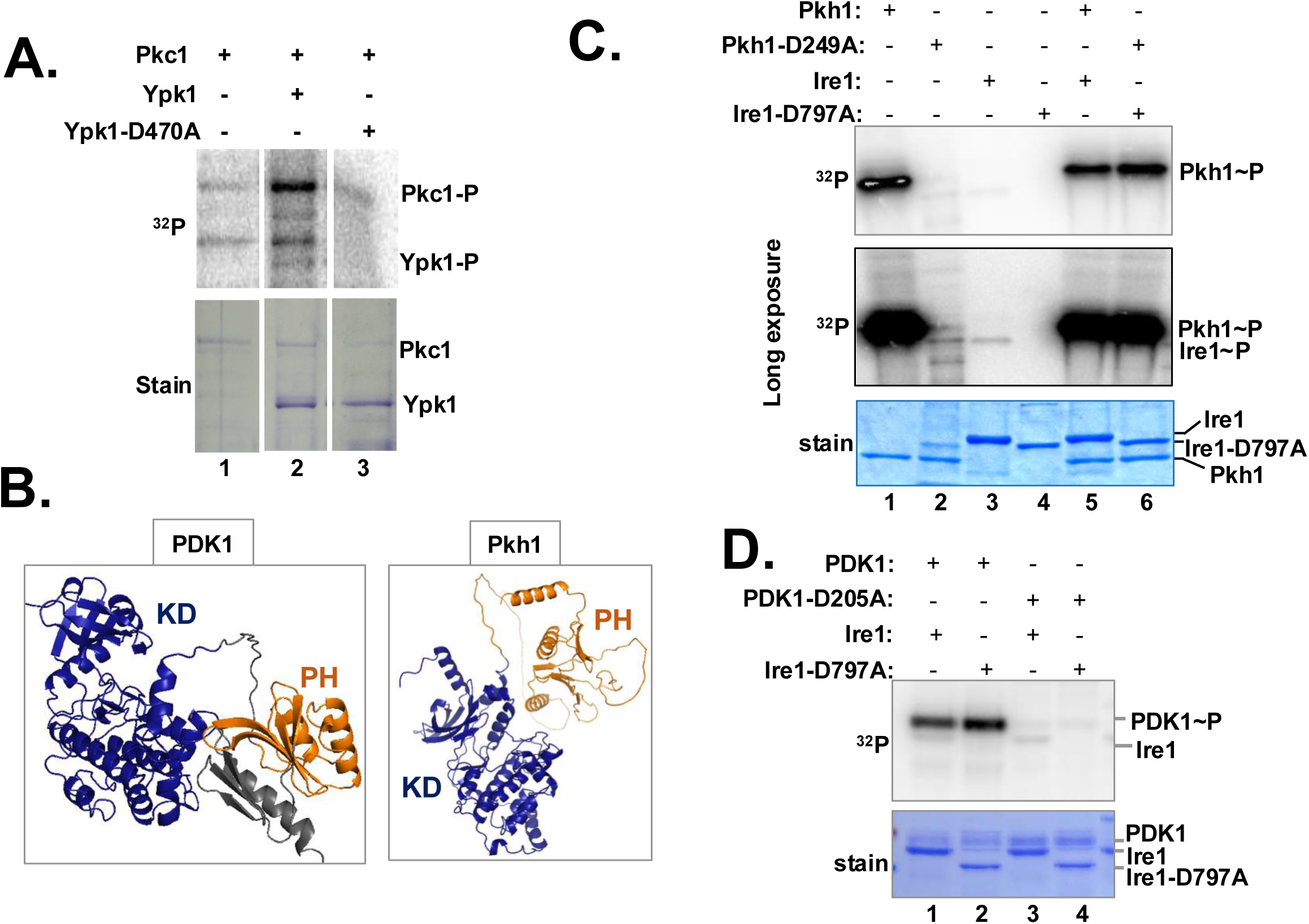
Yeast Pkh1 or its human ortholog PDK1 does not phosphorylate yeast Ire1^cyto^. **(A) Partially purified** Pkc1, Ypk1 or Ypk1-D470A protein mixed in a kinase buffrer in the presence of γ-^32^P-ATP. The reaction mixture was then resolved in an SDS-PAGE gel. The gel was stained, dried and subjected to autoradiography to monitor the ^32^P incorporation in the protein. **(B)** The Alpha-Fold predicted structure of human PDK1 and yeast Pkh1. The kinase domain (KD) and the plecstring homology (PH) domains are colored in blur and orange, respectively. (**C**) & (**CD**) Wild type and kinase-inactive mutant of Pkh1 or PDK1 was partially purified from yeast and subjected to in vitro kinase assays in the presence of γ-^32^P-ATP and recombinant WT and Ire1 kinase-inactive (Ire1-D797A) mutant. The reaction mixture was then resolved in an SDS-PAGE gel. The gel was stained, dried and subjected to autoradiography to monitor the ^32^P incorporation in the protein.

**Supplemental Fig S13:**
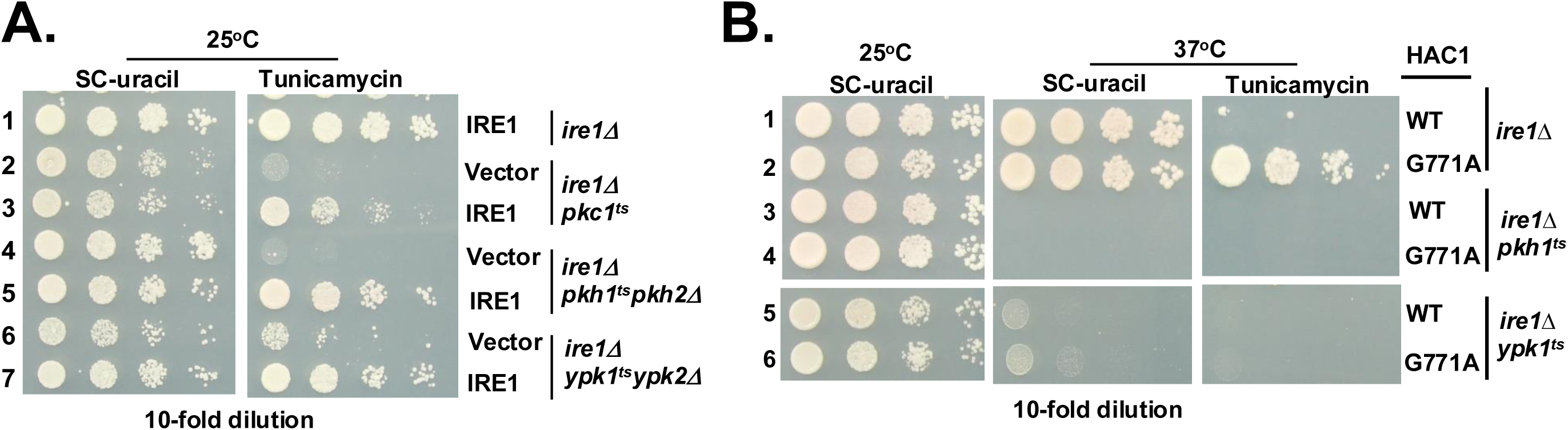
Disruption of *IRE1* gene in *ypk1^ts^ypk2Δ*, *pkh1^ts^pkh2Δ, pkc1^ts^* strain. (A) The *IRE1* gene of the indicated yeast strains was disrupted by the hphMX cassette. The resulting strains were then transformed with a vector plasmid or the same plasmid expressing Ire1. Transformants were then tested for growth on SC-uracil in the presence and absence of tunicamycin. (B) The indicated strains were transformed with a Ura plasmid plasmid expressing WT or G771A mutant of HAC1. Transformants were tested for growth at 25°C and 37°C in the presence and absence of tunicamycin.

**Supplemental Fig S14:**
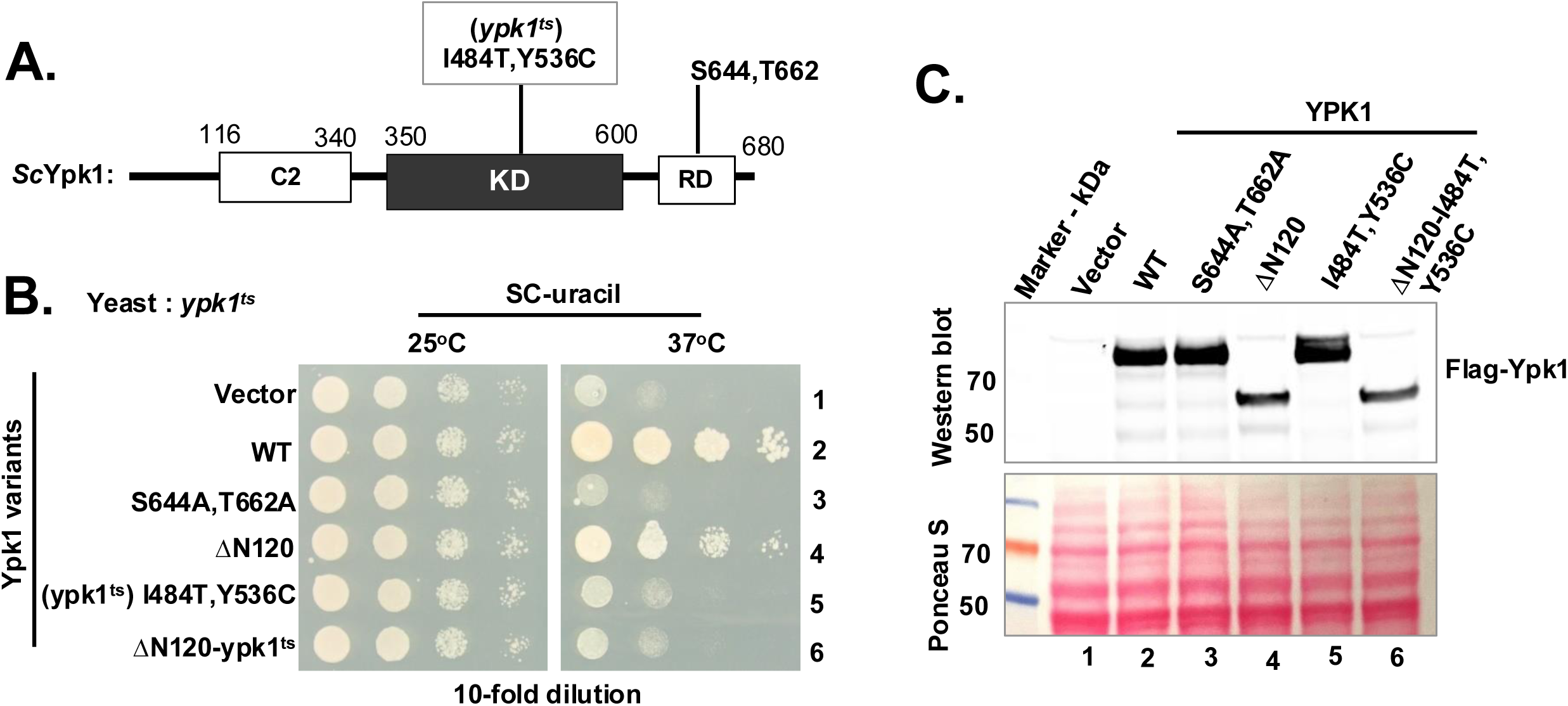
Analysis of Ypk1 mutants. (A) Schematic of *Saccharomyces cerevisiae* Ypk1 (SCYpk1) contain a C2 domain, Kinase domain (KD) and regulatory domain (RD). The numbers indicate the amino acid numbers. Two phospho-acceptor sites (S644 and T662) in the RD and mutations (I484T and Y536C) causing temperature-sensitive phenotypes are shown. (B) The *ypk1^ts^* strain expressing indicated Ypk1 variants were serially diluted and tested for growth at 25°C and 37°C. (C)The whole cell extracts were prepared from strains indicated in (B) and subjected to Western blot analysis using anti Flag antibody to detect Ypk1 protein.

**Supplemental Fig S15:**
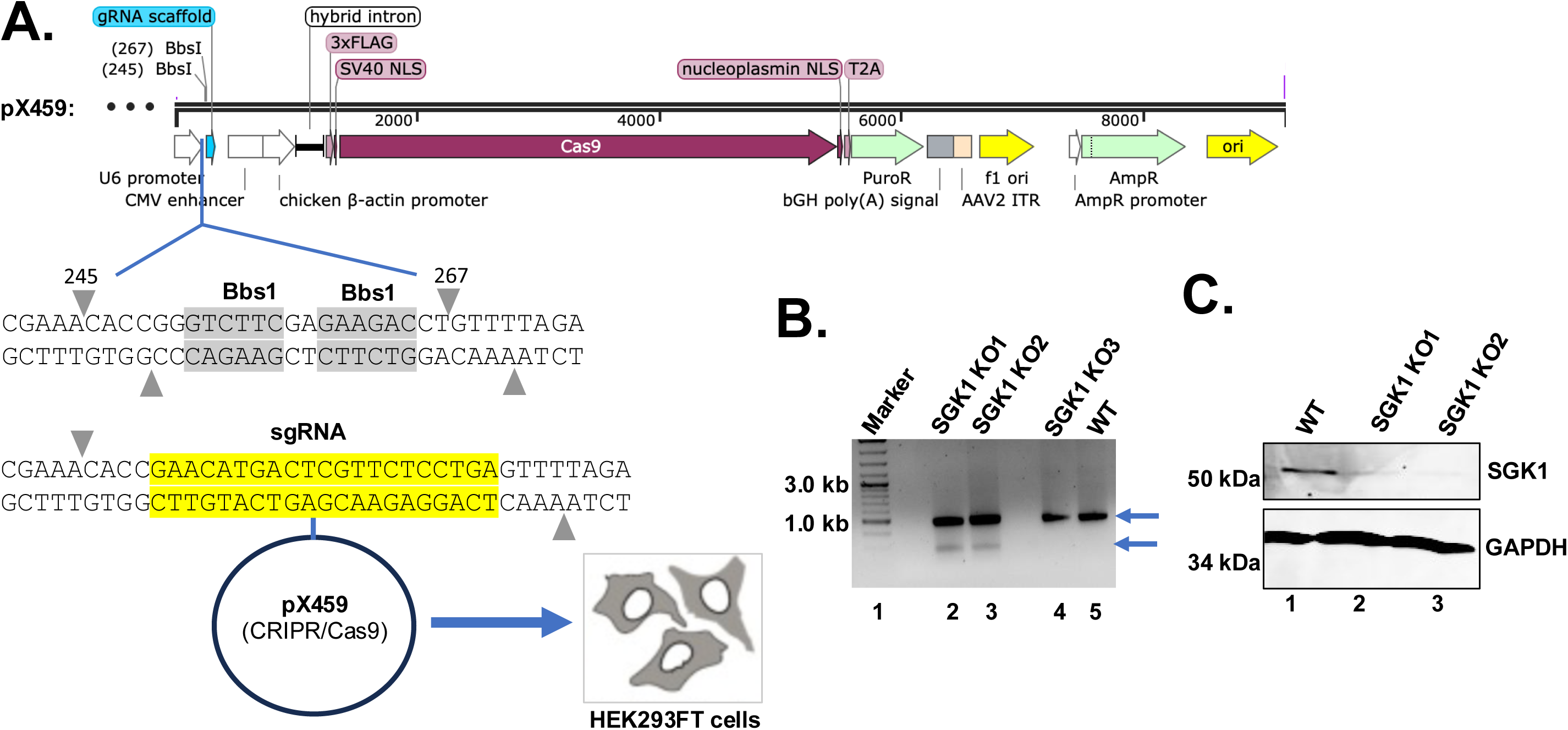
Deletion of SGK1 by CRISPR/Cas9 in HEK293FT cells. (A) The nucleotide sequence encoding the guide RNA targeting the SGK1 locus (sgRNA) was cloned in Bbs1 sites in plasmid pX459. The resulting plasmid was then used to transfect HEK293FT cells. (B) Three putative SGK1 knock out cells (SGK1 KO1, SGK KO2 and SGK1 KO3) were selected. The target DNA sequence from WT and putative knock-out cells was amplified using two gene-specific primers. PCR products were incubated with T7 endonuclease. The digested products were separated in an agarose gel. (C) Whole cell extracts were prepared from WT, SGK1 KO1 and SGK KO2 cells and subjected to Western blot analysis using antibodies raised against SGK1 and GAPDH.

**Supplemental Fig S16:**
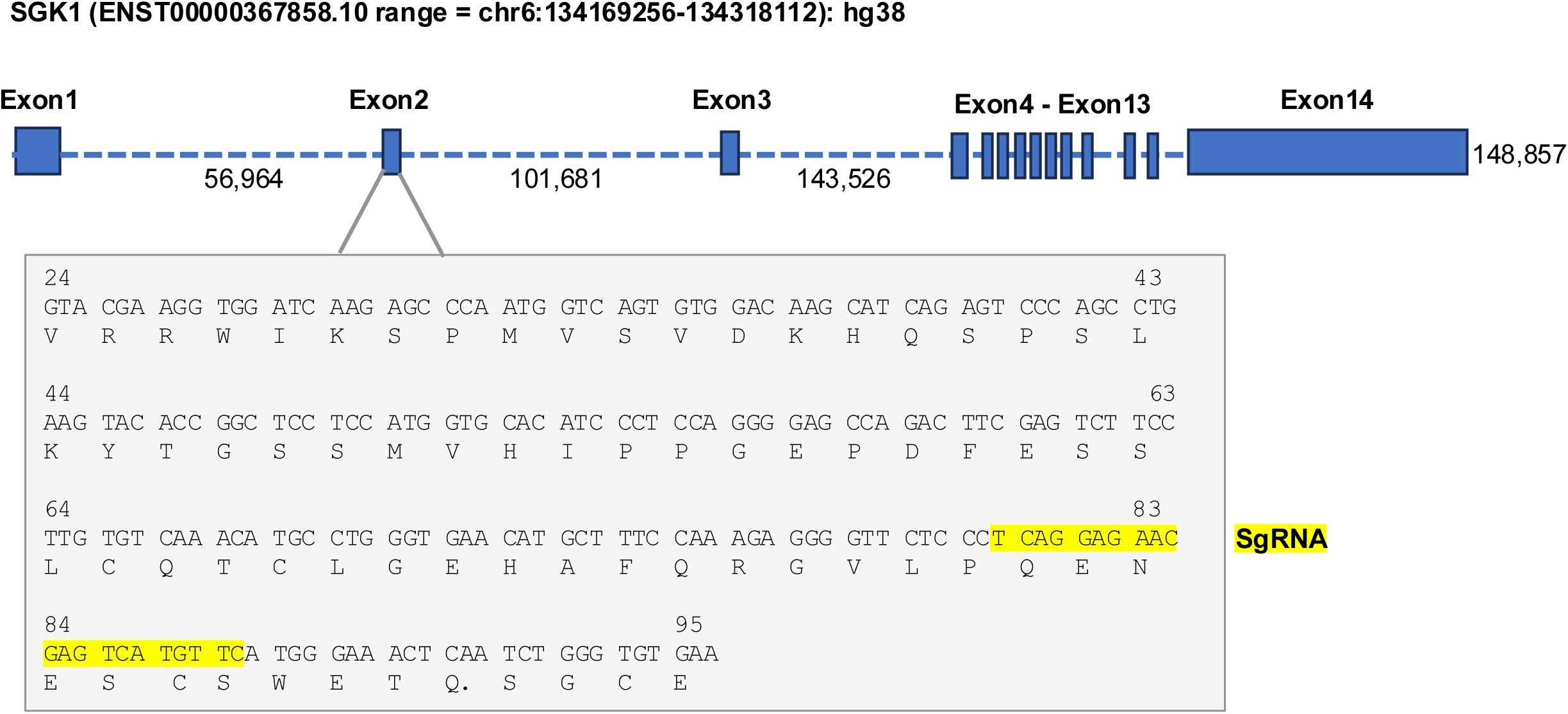
The design of guide RNA (SgRNA) to disrupt SGK1 exons and Introns. The SGK1 gene locus (148, 857 base pairs) comprises of 14 exons (filled boxes) interrupted by introns (dashed lines). Exon2 encodes amino acid residues V24 to E95 shown in a grey box. For gene disruption, the guide RNA, highlighted in yellow, is designed to target the terminal region of exon 2.

**Supplemental Fig S17:**
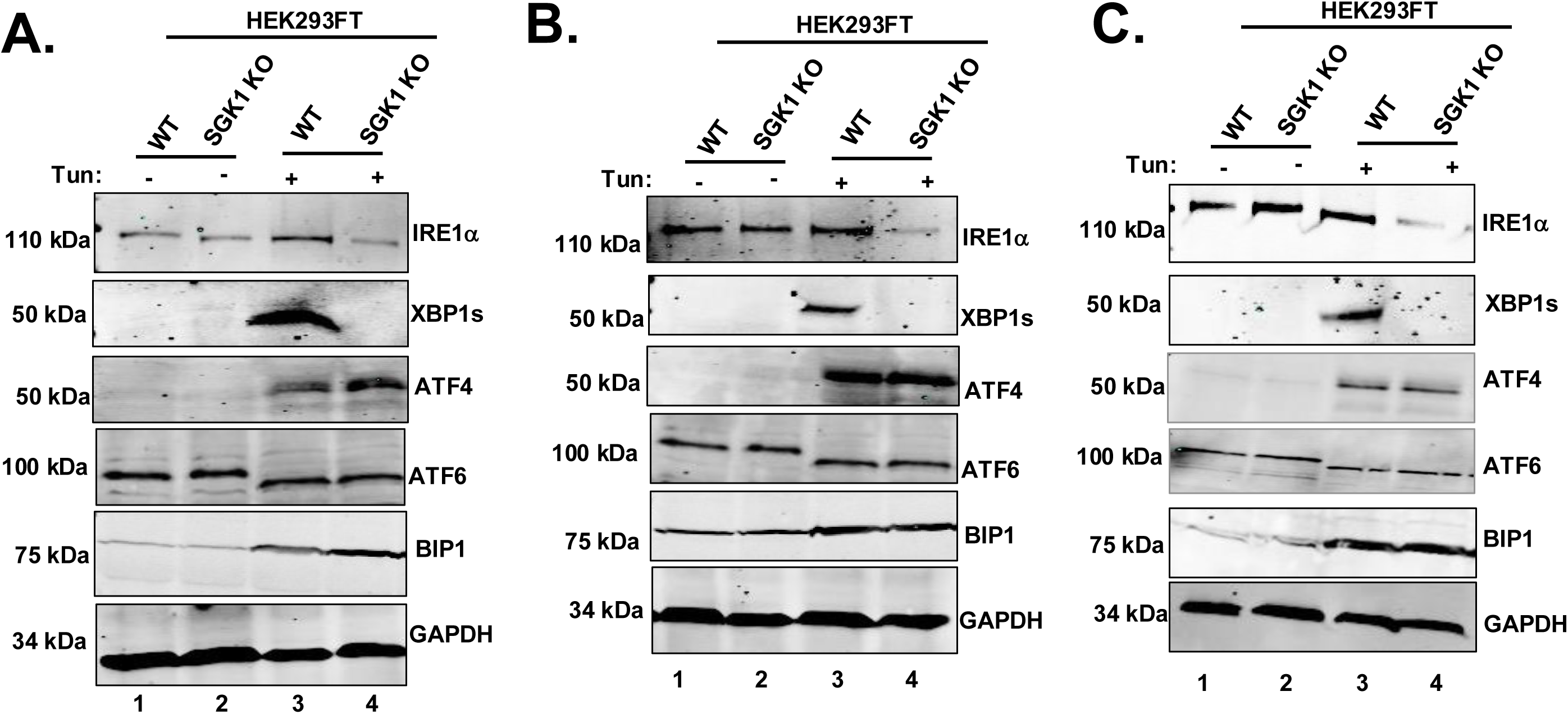
Reduced levels of IRE1 proteins in SGK1-KO cells. Whole cell extracts were prepared from SGK1 knockout cells (SGK1 KO) treated with tunicamycin (Tun, 5μg/ml) and subjected to Western blot analysis by using the indicated antibodies. Three indicated experiments are shown as (A), (B) and (C).

**Supplemental Fig S18:**
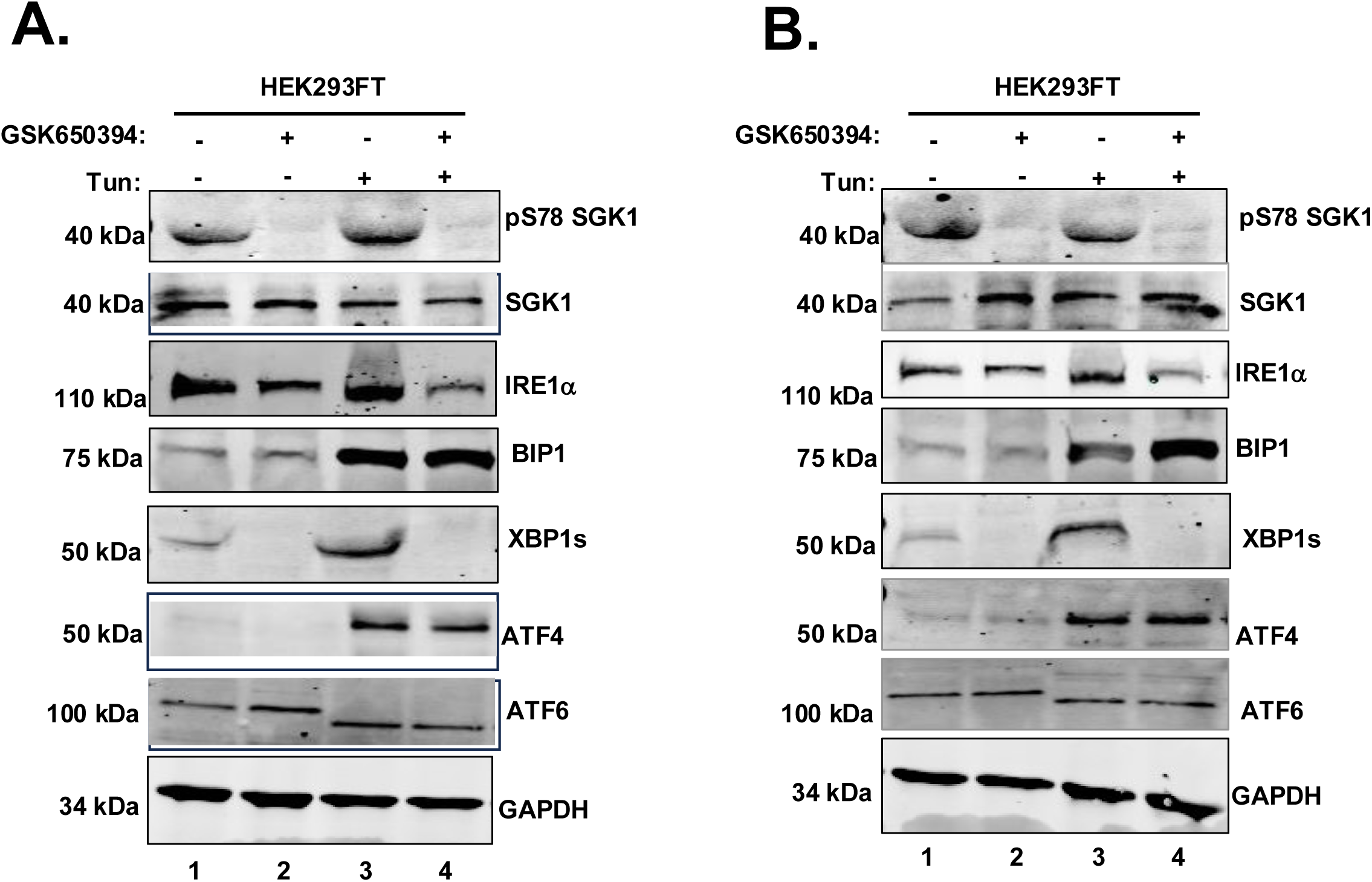
Reduced level of IRE1 in HEK293FT cell treated with SGK1 inhibitor GSK650394. Whole cell extracts were prepared from **HEK293FT** cells treated with and without tunicamycin (Tun, 5μg/ml) and SGK1 inhibitor GSK650394 (5μg/ml) and subjected to Western blot analysis by using the indicated antibodies. Two indicated experiments are shown as (A) and (B).

**Supplemental Fig S19:**
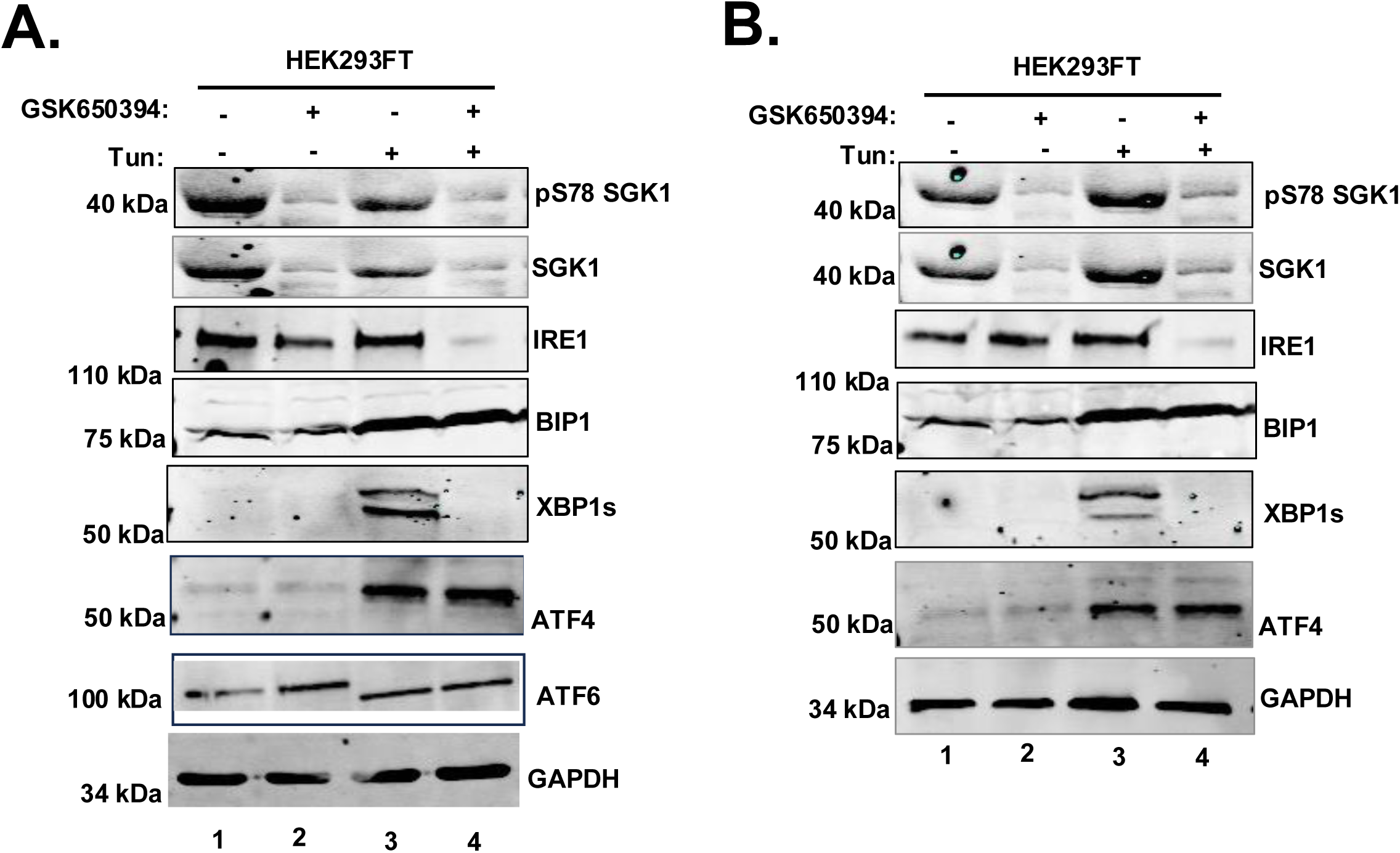
Reduced level of IRE1 protein in ESC WA09 cell treated with SGK1 inhibitor GSK650394. Whole cell extracts were prepared from **ESC WA09** cells treated with and without tunicamycin (Tun, 5μg/ml) and SGK1 inhibitor GSK650394 (5μg/ml) and subjected to Western blot analysis by using the indicated antibodies. Two indicated experiments are shown as (A) and (B).

**Supplemental Fig S20:**
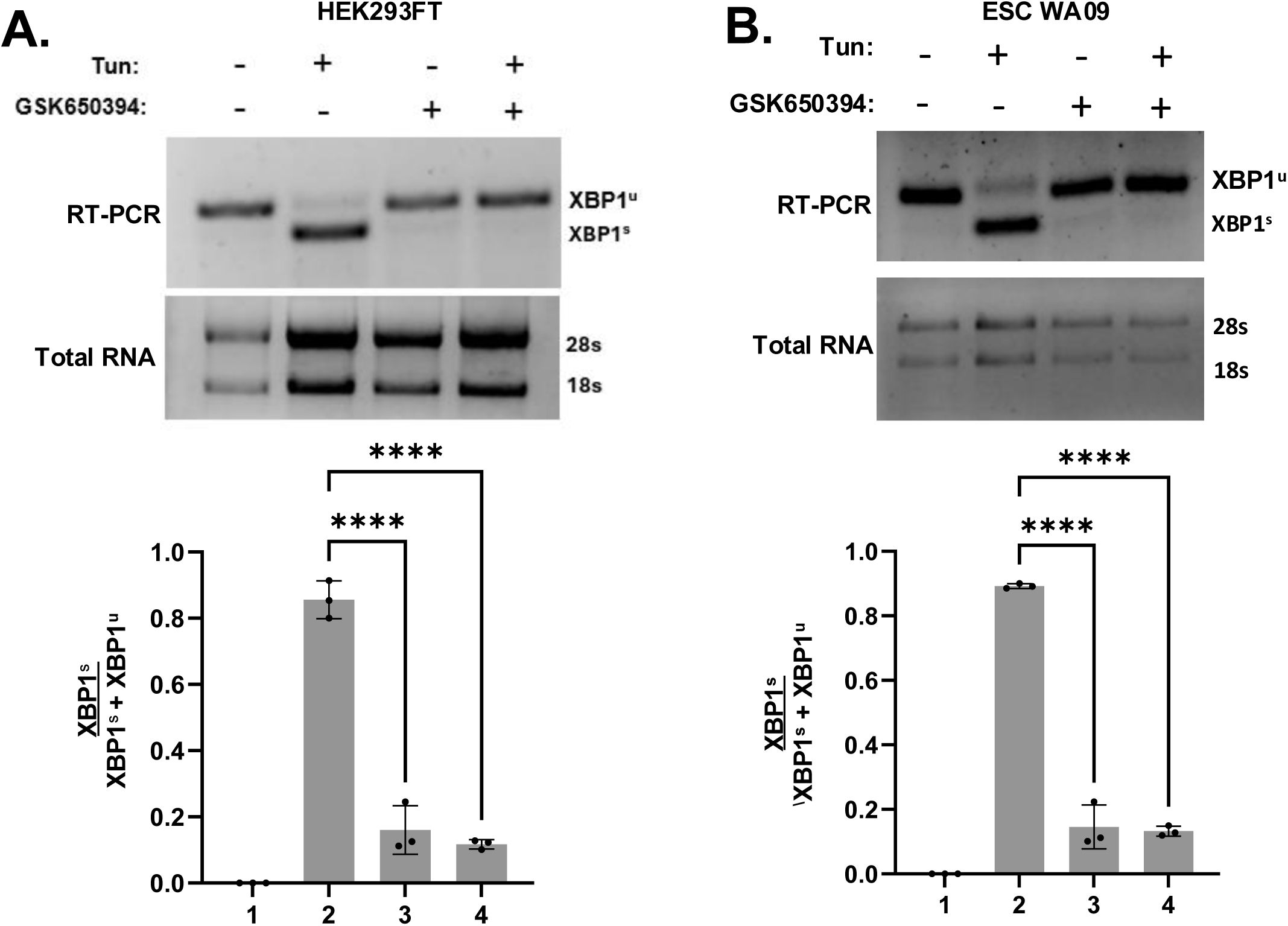
Reduce splicing of XBP1 mRNA in HEK293FT cell treated with SGK1 inhibitor GSK650394. Total RNA was isolated from the HEK293FT (A) and ESC WA09 (B) cells and subjected to RT-PCR to monitor the spliced and un-spliced population of XBP1 mRNAs. (Lower panel). Experiments were repeated three times. The average intensities are shown in a bar diagram (****p-value<0.0001, paired t-test).

## REFERENCES

1. Chen, S., Novick, P., and Ferro-Novick, S. (2013) ER structure and function. Curr Opin Cell Biol 25, 428–433

2. Clapham, D. E. (2007) Calcium signaling. Cell 131, 1047–1058

3. Jacquemyn, J., Cascalho, A., and Goodchild, R. E. (2017) The ins and outs of endoplasmic reticulum-controlled lipid biosynthesis. EMBO Rep 18, 1905–1921

4. Schroder, M. (2008) Endoplasmic reticulum stress responses. Cell Mol Life Sci 65, 862–894

5. Walter, P., and Ron, D. (2011) The unfolded protein response: from stress pathway to homeostatic regulation. Science 334, 1081–1086

6. Ron, D., and Walter, P. (2007) Signal integration in the endoplasmic reticulum unfolded protein response. Nat Rev Mol Cell Bio 8, 519–529

7. Chen, X., Shi, C., He, M., Xiong, S., and Xia, X. (2023) Endoplasmic reticulum stress: molecular mechanism and therapeutic targets. Signal Transduct Target Ther 8, 352

8. Vihervaara, A., Duarte, F. M., and Lis, J. T. (2018) Molecular mechanisms driving transcriptional stress responses. Nat Rev Genet 19, 385–397

9. Wang, X. Z., Harding, H. P., Zhang, Y., Jolicoeur, E. M., Kuroda, M., and Ron, D. (1998) Cloning of mammalian Ire1 reveals diversity in the ER stress responses. EMBO J 17, 5708–5717

10. Tirasophon, W., Welihinda, A. A., and Kaufman, R. J. (1998) A stress response pathway from the endoplasmic reticulum to the nucleus requires a novel bifunctional protein kinase/endoribonuclease (Ire1p) in mammalian cells. Genes Dev 12, 1812–1824

11. Sidrauski, C., and Walter, P. (1997) The transmembrane kinase Ire1p is a site-specific endonuclease that initiates mRNA splicing in the unfolded protein response. Cell 90, 1031–1039

12. Harding, H. P., Zhang, Y., and Ron, D. (1999) Protein translation and folding are coupled by an endoplasmic-reticulum-resident kinase. Nature 397, 271–274

13. Haze, K., Yoshida, H., Yanagi, H., Yura, T., and Mori, K. (1999) Mammalian transcription factor ATF6 is synthesized as a transmembrane protein and activated by proteolysis in response to endoplasmic reticulum stress. Mol Biol Cell 10, 3787–3799

14. Lee, K. P., Dey, M., Neculai, D., Cao, C., Dever, T. E., and Sicheri, F. (2008) Structure of the dual enzyme Ire1 reveals the basis for catalysis and regulation in nonconventional RNA splicing. Cell 132, 89–100

15. Korennykh, A. V., Egea, P. F., Korostelev, A. A., Finer-Moore, J., Zhang, C., Shokat, K. M., Stroud, R. M., and Walter, P. (2009) The unfolded protein response signals through high-order assembly of Ire1. Nature 457, 687–693

16. Prischi, F., Nowak, P. R., Carrara, M., and Ali, M. M. (2014) Phosphoregulation of Ire1 RNase splicing activity. Nat Commun 5, 3554

17. Cox, J. S., and Walter, P. (1996) A novel mechanism for regulating activity of a transcription factor that controls the unfolded protein response. Cell 87, 391–404

18. Ruegsegger, U., Leber, J. H., and Walter, P. (2001) Block of HAC1 mRNA translation by long-range base pairing is released by cytoplasmic splicing upon induction of the unfolded protein response. Cell 107, 103–114

19. Nojima, H., Leem, S. H., Araki, H., Sakai, A., Nakashima, N., Kanaoka, Y., and Ono, Y. (1994) Hac1: a novel yeast bZIP protein binding to the CRE motif is a multicopy suppressor for cdc10 mutant of Schizosaccharomyces pombe. Nucleic Acids Res 22, 5279–5288

20. Chapman, R. E., and Walter, P. (1997) Translational attenuation mediated by an mRNA intron. Curr Biol 7, 850–859

21. Calfon, M., Zeng, H., Urano, F., Till, J. H., Hubbard, S. R., Harding, H. P., Clark, S. G., and Ron, D. (2002) IRE1 couples endoplasmic reticulum load to secretory capacity by processing the XBP-1 mRNA. Nature 415, 92–96

22. Yanagitani, K., Kimata, Y., Kadokura, H., and Kohno, K. (2011) Translational pausing ensures membrane targeting and cytoplasmic splicing of XBP1u mRNA. Science 331, 586–589

23. Sidrauski, C., Cox, J. S., and Walter, P. (1996) tRNA ligase is required for regulated mRNA splicing in the unfolded protein response. Cell 87, 405–413

24. Lu, Y., Liang, F. X., and Wang, X. (2014) A synthetic biology approach identifies the mammalian UPR RNA ligase RtcB. Mol Cell 55, 758–770

25. Travers, K. J., Patil, C. K., Wodicka, L., Lockhart, D. J., Weissman, J. S., and Walter, P. (2000) Functional and genomic analyses reveal an essential coordination between the unfolded protein response and ER-associated degradation. Cell 101, 249–258

26. Harding, H. P., Zhang, Y., Zeng, H., Novoa, I., Lu, P. D., Calfon, M., Sadri, N., Yun, C., Popko, B., Paules, R., Stojdl, D. F., Bell, J. C., Hettmann, T., Leiden, J. M., and Ron, D. (2003) An integrated stress response regulates amino acid metabolism and resistance to oxidative stress. Mol Cell 11, 619–633

27. Lee, A. H., Iwakoshi, N. N., and Glimcher, L. H. (2003) XBP-1 regulates a subset of endoplasmic reticulum resident chaperone genes in the unfolded protein response. Mol Cell Biol 23, 7448–7459

28. Shoulders, M. D., Ryno, L. M., Genereux, J. C., Moresco, J. J., Tu, P. G., Wu, C., Yates, J. R., 3rd, Su, A. I., Kelly, J. W., and Wiseman, R. L. (2013) Stress-independent activation of XBP1s and/or ATF6 reveals three functionally diverse ER proteostasis environments. Cell Rep 3, 1279–1292

29. Van Dalfsen, K. M., Hodapp, S., Keskin, A., Otto, G. M., Berdan, C. A., Higdon, A., Cheunkarndee, T., Nomura, D. K., Jovanovic, M., and Brar, G. A. (2018) Global Proteome Remodeling during ER Stress Involves Hac1-Driven Expression of Long Undecoded Transcript Isoforms. Dev Cell 46, 219–235 e218

30. Chen, Y., Feldman, D. E., Deng, C., Brown, J. A., De Giacomo, A. F., Gaw, A. F., Shi, G., Le, Q. T., Brown, J. M., and Koong, A. C. (2005) Identification of mitogen-activated protein kinase signaling pathways that confer resistance to endoplasmic reticulum stress in Saccharomyces cerevisiae. Mol Cancer Res 3, 669–677

31. Chakraborty, A., Chakrabarty, S., Uppala, J. K., Mayer, K. A., Evans, A. J., George, J., Ghosh, C., Dey, R., Ohikhuare, F., Mirza, S., Nguyen, A. P. T., Gouignard, N., Chaluvally-Raghavan, P., and Dey, M. (2026) The transcription factor Rlm1 couples the MAPK Slt2/ERK1 pathway to the IRE1-driven unfolded protein response. Commun Biol 9

32. Uppala, J. K., Bhattacharjee, S., and Dey, M. (2021) Vps34 and TOR Kinases Coordinate HAC1 mRNA Translation in the Presence or Absence of Ire1-Dependent Splicing. Mol Cell Biol 41, e0066220

33. Pastor-Flores, D., Ferrer-Dalmau, J., Bahi, A., Boleda, M., Biondi, R. M., and Casamayor, A. (2015) Depletion of yeast PDK1 orthologs triggers a stress-like transcriptional response. BMC Genomics 16, 719

34. Casamayor, A., Torrance, P. D., Kobayashi, T., Thorner, J., and Alessi, D. R. (1999) Functional counterparts of mammalian protein kinases PDK1 and SGK in budding yeast. Curr Biol 9, 186–197

35. Roelants, F. M., Breslow, D. K., Muir, A., Weissman, J. S., and Thorner, J. (2011) Protein kinase Ypk1 phosphorylates regulatory proteins Orm1 and Orm2 to control sphingolipid homeostasis in Saccharomyces cerevisiae. Proc Natl Acad Sci U S A 108, 19222–19227

36. Li, Z., Vizeacoumar, F. J., Bahr, S., Li, J., Warringer, J., Vizeacoumar, F. S., Min, R., Vandersluis, B., Bellay, J., Devit, M., Fleming, J. A., Stephens, A., Haase, J., Lin, Z. Y., Baryshnikova, A., Lu, H., Yan, Z., Jin, K., Barker, S., Datti, A., Giaever, G., Nislow, C., Bulawa, C., Myers, C. L., Costanzo, M., Gingras, A. C., Zhang, Z., Blomberg, A., Bloom, K., Andrews, B., and Boone, C. (2011) Systematic exploration of essential yeast gene function with temperature-sensitive mutants. Nat Biotechnol 29, 361–367

37. Schmidt, A., Kunz, J., and Hall, M. N. (1996) TOR2 is required for organization of the actin cytoskeleton in yeast. Proc Natl Acad Sci U S A 93, 13780–13785

38. Luo, G., Gruhler, A., Liu, Y., Jensen, O. N., and Dickson, R. C. (2008) The sphingolipid long-chain base-Pkh1/2-Ypk1/2 signaling pathway regulates eisosome assembly and turnover. J Biol Chem 283, 10433–10444

39. Roelants, F. M., Torrance, P. D., Bezman, N., and Thorner, J. (2002) Pkh1 and Pkh2 differentially phosphorylate and activate Ypk1 and Ykr2 and define protein kinase modules required for maintenance of cell wall integrity. Mol Biol Cell 13, 3005–3028

40. Panek, H. R., Stepp, J. D., Engle, H. M., Marks, K. M., Tan, P. K., Lemmon, S. K., and Robinson, L. C. (1997) Suppressors of YCK-encoded yeast casein kinase 1 deficiency define the four subunits of a novel clathrin AP-like complex. EMBO J 16, 4194–4204

41. Hartwell, L. H., Culotti, J., and Reid, B. (1970) Genetic control of the cell-division cycle in yeast. I. Detection of mutants. Proc Natl Acad Sci U S A 66, 352–359

42. Kunz, J., Henriquez, R., Schneider, U., Deuter-Reinhard, M., Movva, N. R., and Hall, M. N. (1993) Target of rapamycin in yeast, TOR2, is an essential phosphatidylinositol kinase homolog required for G1 progression. Cell 73, 585–596

43. Levin, D. E., Fields, F. O., Kunisawa, R., Bishop, J. M., and Thorner, J. (1990) A candidate protein kinase C gene, PKC1, is required for the S. cerevisiae cell cycle. Cell 62, 213–224

44. Geerlings, T. H., Faber, A. W., Bister, M. D., Vos, J. C., and Raue, H. A. (2003) Rio2p, an evolutionarily conserved, low abundant protein kinase essential for processing of 20 S Pre-rRNA in Saccharomyces cerevisiae. J Biol Chem 278, 22537–22545

45. deHart, A. K., Schnell, J. D., Allen, D. A., and Hicke, L. (2002) The conserved Pkh-Ypk kinase cascade is required for endocytosis in yeast. J Cell Biol 156, 241–248

46. Niles, B. J., Joslin, A. C., Fresques, T., and Powers, T. (2014) TOR complex 2-Ypk1 signaling maintains sphingolipid homeostasis by sensing and regulating ROS accumulation. Cell Rep 6, 541–552

47. Leskoske, K. L., Roelants, F. M., Emmerstorfer-Augustin, A., Augustin, C. M., Si, E. P., Hill, J. M., and Thorner, J. (2018) Phosphorylation by the stress-activated MAPK Slt2 down-regulates the yeast TOR complex 2. Genes Dev 32, 1576–1590

48. Niles, B. J., Mogri, H., Hill, A., Vlahakis, A., and Powers, T. (2012) Plasma membrane recruitment and activation of the AGC kinase Ypk1 is mediated by target of rapamycin complex 2 (TORC2) and its effector proteins Slm1 and Slm2. Proc Natl Acad Sci U S A 109, 1536–1541

49. Roelants, F. M., Baltz, A. G., Trott, A. E., Fereres, S., and Thorner, J. (2010) A protein kinase network regulates the function of aminophospholipid flippases. Proc Natl Acad Sci U S A 107, 34–39

50. Lee, Y. J., Jeschke, G. R., Roelants, F. M., Thorner, J., and Turk, B. E. (2012) Reciprocal phosphorylation of yeast glycerol-3-phosphate dehydrogenases in adaptation to distinct types of stress. Mol Cell Biol 32, 4705–4717

51. Han, S., Lone, M. A., Schneiter, R., and Chang, A. (2010) Orm1 and Orm2 are conserved endoplasmic reticulum membrane proteins regulating lipid homeostasis and protein quality control. Proc Natl Acad Sci U S A 107, 5851–5856

52. Nomoto, S., Watanabe, Y., Ninomiya-Tsuji, J., Yang, L. X., Nagai, Y., Kiuchi, K., Hagiwara, M., Hidaka, H., Matsumoto, K., and Irie, K. (1997) Functional analyses of mammalian protein kinase C isozymes in budding yeast and mammalian fibroblasts. Genes Cells 2, 601–614

53. Inagaki, M., Schmelzle, T., Yamaguchi, K., Irie, K., Hall, M. N., and Matsumoto, K. (1999) PDK1 homologs activate the Pkc1-mitogen-activated protein kinase pathway in yeast. Mol Cell Biol 19, 8344–8352

54. Lee, K. S., and Levin, D. E. (1992) Dominant mutations in a gene encoding a putative protein kinase (BCK1) bypass the requirement for a Saccharomyces cerevisiae protein kinase C homolog. Mol Cell Biol 12, 172–182

55. Turk, B. E. (2008) Understanding and exploiting substrate recognition by protein kinases. Curr Opin Chem Biol 12, 4–10

56. Mannan, M. A., Shadrick, W. R., Biener, G., Shin, B. S., Anshu, A., Raicu, V., Frick, D. N., and Dey, M. (2013) An ire1-phk1 chimera reveals a dispensable role of autokinase activity in endoplasmic reticulum stress response. J Mol Biol 425, 2083–2099

57. Uppala, J. K., Sathe, L., Chakraborty, A., Bhattacharjee, S., Pulvino, A. T., and Dey, M. (2022) The cap-proximal RNA secondary structure inhibits preinitiation complex formation on HAC1 mRNA. J Biol Chem 298, 101648

58. Surlow, B. A., Cooley, B. M., Needham, P. G., Brodsky, J. L., and Patton-Vogt, J. (2014) Loss of Ypk1, the yeast homolog to the human serum- and glucocorticoid-induced protein kinase, accelerates phospholipase B1-mediated phosphatidylcholine deacylation. J Biol Chem 289, 31591–31604

59. Nalefski, E. A., and Falke, J. J. (1996) The C2 domain calcium-binding motif: structural and functional diversity. Protein Sci 5, 2375–2390

60. Larsen, A. H., and Sansom, M. S. P. (2021) Binding of Ca(2+)-independent C2 domains to lipid membranes: A multi-scale molecular dynamics study. Structure 29, 1200–1213 e1202

61. Varadi, M., Anyango, S., Deshpande, M., Nair, S., Natassia, C., Yordanova, G., Yuan, D., Stroe, O., Wood, G., Laydon, A., Zidek, A., Green, T., Tunyasuvunakool, K., Petersen, S., Jumper, J., Clancy, E., Green, R., Vora, A., Lutfi, M., Figurnov, M., Cowie, A., Hobbs, N., Kohli, P., Kleywegt, G., Birney, E., Hassabis, D., and Velankar, S. (2022) AlphaFold Protein Structure Database: massively expanding the structural coverage of protein-sequence space with high-accuracy models. Nucleic Acids Res 50, D439–D444

62. Anacker, C., Cattaneo, A., Musaelyan, K., Zunszain, P. A., Horowitz, M., Molteni, R., Luoni, A., Calabrese, F., Tansey, K., Gennarelli, M., Thuret, S., Price, J., Uher, R., Riva, M. A., and Pariante, C. M. (2013) Role for the kinase SGK1 in stress, depression, and glucocorticoid effects on hippocampal neurogenesis. Proc Natl Acad Sci U S A 110, 8708–8713

63. Ye, J., Rawson, R. B., Komuro, R., Chen, X., Dave, U. P., Prywes, R., Brown, M. S., and Goldstein, J. L. (2000) ER stress induces cleavage of membrane-bound ATF6 by the same proteases that process SREBPs. Mol Cell 6, 1355–1364

64. Cox, J. S., Shamu, C. E., and Walter, P. (1993) Transcriptional Induction of Genes Encoding Endoplasmic-Reticulum Resident Proteins Requires a Transmembrane Protein-Kinase. Cell 73, 1197–1206

65. Peschek, J., Acosta-Alvear, D., Mendez, A. S., and Walter, P. (2015) A conformational RNA zipper promotes intron ejection during non-conventional XBP1 mRNA splicing. EMBO Rep 16, 1688–1698

66. Ruegsegger, U., Leber, J. H., and Walter, P. (2001) Block of HAC1 mRNA translation by long-range base pairing is released by cytoplasmic splicing upon induction of the unfolded protein response. Cell 107, 103–114

67. Hinnebusch, A. G. (2005) Translational regulation of GCN4 and the general amino acid control of yeast. Annual review of microbiology 59, 407–450

68. Harding, H. P., Novoa, I., Zhang, Y., Zeng, H., Wek, R., Schapira, M., and Ron, D. (2000) Regulated translation initiation controls stress-induced gene expression in mammalian cells. Mol Cell 6, 1099–1108

69. Kimata, Y., Kimata, Y. I., Shimizu, Y., Abe, H., Farcasanu, I. C., Takeuchi, M., Rose, M. D., and Kohno, K. (2003) Genetic evidence for a role of BiP/Kar2 that regulates Ire1 in response to accumulation of unfolded proteins. Mol Biol Cell 14, 2559–2569

70. Kohno, K., Normington, K., Sambrook, J., Gething, M. J., and Mori, K. (1993) The promoter region of the yeast KAR2 (BiP) gene contains a regulatory domain that responds to the presence of unfolded proteins in the endoplasmic reticulum. Mol Cell Biol 13, 877–890

71. Carrara, M., Prischi, F., Nowak, P. R., Kopp, M. C., and Ali, M. M. (2015) Noncanonical binding of BiP ATPase domain to Ire1 and Perk is dissociated by unfolded protein CH1 to initiate ER stress signaling. eLife 4

72. Wooden, S. K., and Lee, A. S. (1992) Comparison of the genomic organizations of the rat grp78 and hsc73 gene and their evolutionary implications. DNA Seq 3, 41–48

73. Lee, A. S. (2005) The ER chaperone and signaling regulator GRP78/BiP as a monitor of endoplasmic reticulum stress. Methods 35, 373–381

74. Breitkreutz, A., Choi, H., Sharom, J. R., Boucher, L., Neduva, V., Larsen, B., Lin, Z. Y., Breitkreutz, B. J., Stark, C., Liu, G., Ahn, J., Dewar-Darch, D., Reguly, T., Tang, X., Almeida, R., Qin, Z. S., Pawson, T., Gingras, A. C., Nesvizhskii, A. I., and Tyers, M. (2010) A global protein kinase and phosphatase interaction network in yeast. Science 328, 1043–1046

75. Levin, D. E. (2011) Regulation of cell wall biogenesis in Saccharomyces cerevisiae: the cell wall integrity signaling pathway. Genetics 189, 1145–1175

76. Gustin, M. C., Albertyn, J., Alexander, M., and Davenport, K. (1998) MAP kinase pathways in the yeast Saccharomyces cerevisiae. Microbiol Mol Biol Rev 62, 1264–1300

77. Irie, K., Takase, M., Lee, K. S., Levin, D. E., Araki, H., Matsumoto, K., and Oshima, Y. (1993) MKK1 and MKK2, which encode Saccharomyces cerevisiae mitogen-activated protein kinase-kinase homologs, function in the pathway mediated by protein kinase C. Mol Cell Biol 13, 3076–3083

78. Zarzov, P., Mazzoni, C., and Mann, C. (1996) The SLT2(MPK1) MAP kinase is activated during periods of polarized cell growth in yeast. EMBO J 15, 83–91

79. Pearce, L. R., Komander, D., and Alessi, D. R. (2010) The nuts and bolts of AGC protein kinases. Nat Rev Mol Cell Biol 11, 9–22

80. Jang, H., Park, Y., and Jang, J. (2022) Serum and glucocorticoid-regulated kinase 1: Structure, biological functions, and its inhibitors. Front Pharmacol 13, 1036844

81. Sakuma, T., Nishikawa, A., Kume, S., Chayama, K., and Yamamoto, T. (2014) Multiplex genome engineering in human cells using all-in-one CRISPR/Cas9 vector system. Sci Rep 4, 5400

82. Schneider, C. A., Rasband, W. S., and Eliceiri, K. W. (2012) NIH Image to ImageJ: 25 years of image analysis. Nat Methods 9, 671–675

